# Chromosome length is not the sole determinant of sexually dimorphic crossover rates during mammalian meiosis: Insights from genetically diverse mouse strains

**DOI:** 10.64898/2025.12.19.695521

**Authors:** Tegan S. Horan, Anna Wood, Stephanie Tanis, Carme Peirau Gabarrell, Paula E. Cohen

**Affiliations:** Department of Biomedical Sciences, Cornell University, Ithaca, NY 14853; Cornell Reproductive Sciences Center, Cornell University, Ithaca, NY 14853

**Keywords:** Crossover, Meiotic Recombination, Crossover Interference, Sexual Dimorphism

## Abstract

Meiotic recombination generates crossovers (COs), reciprocal exchanges between homologous chromosomes critical for accurate chromosome segregation. Inappropriate CO frequency and distribution drive aneuploidy in human oocytes, with error rates up to 10-fold higher than in sperm despite females exhibiting higher CO frequencies. COs form in the context of the proteinaceous synaptonemal complex (SC) that tethers homologs during prophase I. SC length strongly correlates with CO number, and sexual dimorphism in recombination has long been attributed to longer SCs in females. However, this model is challenged by wild-derived PWD mice in which males consistently generate more COs despite having shorter SCs. Here, we exploit natural genetic variation among inbred mouse strains to dissect the structural and regulatory basis of sexually dimorphic CO regulation. Using cytological markers of SC assembly (SYCP3), recombination progression (RAD51, MSH4), class I CO designation (HEI10, MLH1/MLH3), and chiasmata, we show that SC length is not the sole predictor of CO number. PWD males exhibit stronger CO interference and higher CO number than females, despite reduced SC length. Notably, females show reduced efficiency in designating recombination intermediate to become COs, whereas PWD males display exceptional proficiency. Unexpectedly, although class II COs are rare, they play a disproportionate role in ensuring that every chromosome pair receives at least one CO, thereby safeguarding against aneuploidy. Together, these findings challenge the prevailing view that SC length is the primary determinant of sexually dimorphic CO rates and instead highlight sex-specific regulation of CO designation and pathway usage as key drivers of recombination outcomes.

## Introduction

Germ cells have a formidable task during meiosis: they must halve their chromosome number through two cell divisions without intervening DNA replication to ensure each resulting gamete (sperm or egg) inherits exactly one copy of each chromosome. To accomplish this, the two parental chromosomes (homologs) must recognize each other and remain paired throughout prophase I via two interconnected processes. First, a protein structure, the synaptonemal complex (SC), forms between and tethers homologs together. Second, crossing over creates reciprocal DNA exchanges (crossovers, COs) that physically link homologs. These COs serve multiple functions: they maintain pairing through metaphase I, facilitate proper orientation and spindle attachment, and ensure optimal tension forms across the centromeres/kinetochores for homologs to be pulled to opposite poles^1–4^. Crossing over is tightly regulated to ensure each homologous pair receives at minimum one obligate CO (assurance), without which chromosomes segregate stochastically and frequently produce chromosomally atypical (aneuploid) gametes^5,6^. In mammals, COs are equally critical during spermatogenesis and oogenesis, and both processes utilize the same core DNA repair machinery to produce COs from an abundant pool of precursor events^7^. However, CO regulation shows considerable sexual dimorphism and produces male-female differences in both the number of COs (heterochiasmy) and their distribution along the chromosome^8,9^. Importantly in humans, these sex differences contribute to a high incidence of aneuploidy (roughly 10% of conceptuses), most originating from errors in maternal meiosis; as many as 20 - 80% of eggs (depending on age) contrasted with 2.5 - 7% of sperm have segregation errors^10–13^. Achiasmy (homologous chromosomes that fail to make the obligate CO) is the best-characterized source of aneuploidy in humans^10,14^. Recent studies have shown that achiasmy is both common in human oocytes and its incidence is higher in oocytes with reduced global CO rates^15^. Importantly, growing evidence indicates sub-optimal CO positioning (e.g., spaced too close together or too near the centromere or telomere) also strongly associate with maternal segregation errors ^10,12,16^.

Much of our understanding of the molecular regulation of CO processing comes from biochemical studies of *S. cerevisiae*, with confirmation of shared regulatory events in many meiotic species^6,17,18^. In all cases, SPO11 and its accessory proteins induce programmed double-strand breaks (DSBs) throughout the genome at the onset of meiosis^19–21^. Of more than 200 breaks made in mice and humans, only ∼10% are resolved as COs^6,18^. The remaining DSBs become non-crossovers (NCOs) via synthesis-dependent strand annealing (SDSA) or the helicase-mediated dissolution of repair intermediates^22–25^. Of the DSBs that resolve as COs, nearly all (90-95%) form via the class I pathway mediated by DNA mismatch repair (MMR) proteins: During CO licensing, MutSγ (MSH4-MSH5) binds and stabilizes repair intermediates, and a subset of these undergo designation and are resolved by MutLγ (MLH1-MLH3) as COs ^22,26–32^. Most of the remaining 5-10% of COs are the result of the minor class II pathway and mediated by the structure-selective nuclease (SSN), MUS81-EME1^6,33,34^.

Both the number and the distribution of COs show striking sex differences: In most mammals, females have more COs than males, and males have more distally positioned COs than females^7,9,35,36^. Meiotic recombination occurs specifically within the context of the chromosome axis. The number and spacing of class I COs along the axis are restricted by interference, a process that leads to the inhibition of two adjacent class I COs from forming within a close physical distance that is mediated in part by the SC^37–42^. It is perhaps not surprising, therefore, that SC length also exhibits sexual dimorphism. Most female mammals have longer SCs than males, and a strong covariation between CO frequency and SC length is well established ^35,43–47^. In humans and mice, sex differences in interference distance tend to be larger when measured in genomic distance (megabases of DNA); however inter-CO spacing is largely comparable at the level of micron length along the SC, leading to the long-held understanding that sex differences in CO frequency are attributable to sex differences in SC length^39,44,48–50^.

Most of what is known about the regulation of meiotic recombination in mammals comes from studies of mice, especially the C57BL/6J strain. Further, due to the difficulty inherent to female prophase I studies — these events arising during embryogenesis in females — most mammalian data come from investigations of spermatogenesis. Sex and strain differences in CO rates have previously been defined, but few studies have characterized strain differences in DSB formation or CO processing, and these observations have been confined only to males^51–56^. Recent studies have shown that many wild-derived mouse strains from the *Mus musculus musculus* subspecies exhibit reversed patterns of heterochiasmy whereby males from the PWD/PhJ (PWD) strain make more class I COs than females, but curiously retain a shorter SC length like males from other strains^53,57^. These unique qualities make PWD mice powerful tools for examining sex differences in the regulation of crossing over and the significance of chromosome axis length in heterochiasmy. To better understand the importance of chromosome axis length on sexually dimorphic CO frequency and patterning, we conducted an in-depth cytological analysis of CO distribution and synaptonemal complex length in five inbred mouse strains. We selected strains in which male class I CO frequencies (as measured by MLH1 foci) were low (DBA/2J [DBA]^58^, CAST/EiJ [CAST]^51,52,54^), intermediate (C57BL/6J [B6]^51,52,59^, 129S1/SvImJ [129S1]^60^), or high (PWD/PhJ [PWD]^53,54^). This panel of strains was also chosen to capture a high degree of genetic diversity, comprising three subspecies (*M. m. domesticus* [DBA, 129S1, B6], *M. m. musculus* [PWD], and *M. m. castaneus* [CAST]), and four of the seven phylogenetic groups identified by Petkov et al.^61^ We also performed a thorough analysis of sex differences in the trajectory of recombination from DSBs through COs in PWD and B6 mice. Our work reveals critical multifactorial sex and strain differences in CO regulation that disrupt the long-held understanding that sex differences in SC length are the predominant driver of mammalian heterochiasmy.

## Results

### Sex and strain differences in class I and II crossovers

Prior comparisons of inter-strain sex differences in mouse CO frequency have used MLH1 foci as their only metric of crossovers, leaving critical questions about variation in the frequency of alternative CO pathways (e.g., the MUS81-EME1 mediated class II pathway) unanswered. To validate these previous findings and expand on them to examine total CO numbers, we used both indirect immunofluorescence (IF) staining of MLH1 to assess class I COs in pachynema in both adult spermatocytes and fetal oocytes (Fig. 1A-D; Fig. S1A-B, E-F, I-J, M-N, Q-R).

**Fig. 1.**
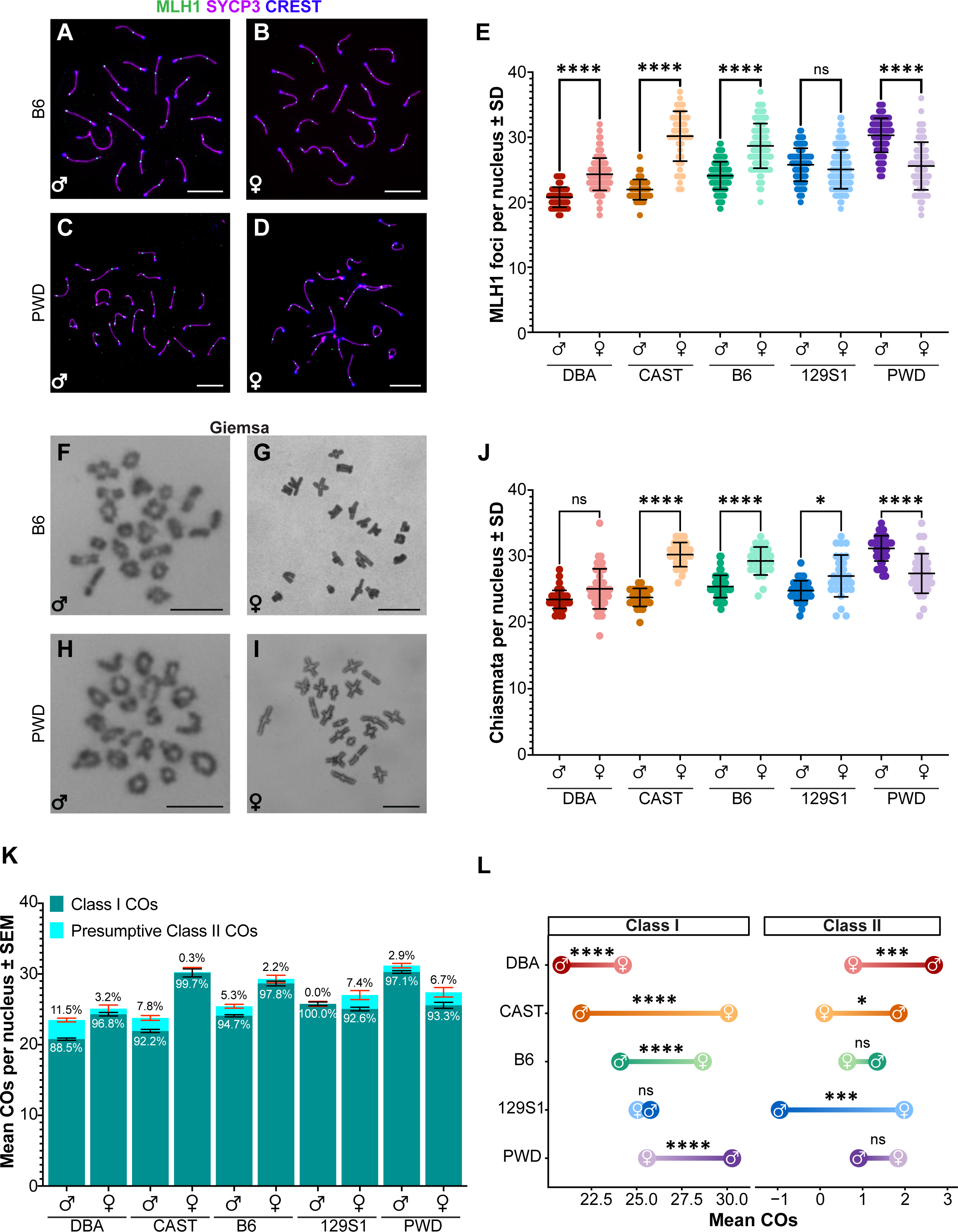
P**a**tterns **of sexual dimorphism in class I and II crossovers are inconsistent among diverse mouse strains.** Representative immunofluorescence images of MLH1 (green, class I COs), SYCP3 (violet, chromosome axes), and centromeres (blue) in spermatocyte (A, C) and oocyte (B, D) chromosome spreads from B6 and PWD mice, respectively. (E) Significant sex differences in mean MLH1 foci number were observed in DBA (red, t = 11.6), CAST (orange, t = 13.4), B6 (green, t = 12.4), and PWD (purple, t = 10.2), but not 129S1 (blue, t = 1.9) mice. Representative Giemsa-stained chiasmata spreads from diakinesis/metaphase I spermatocytes (F, H) and oocytes (G, I) from B6 and PWD mice, respectively. (J) Significant sex differences in mean chiasmata number were observed in CAST (t = 14.8), B6 (t = 8.3), 129S1 (t = 3.5), and PWD (t = 6.6) but not DBA (t = 3.2) mice. (K) Presumptive class II CO numbers (light teal) were calculated by subtracting mean MLH1 foci (dark teal) from mean chiasmata for each sex and strain; class II CO standard error (red bars) was calculated by error propagation assuming independence between measures. (L) Dumbbell plot of sex differences in CO pathway usage shows that the sex with more class I COs consistently has fewer class II COs, with significant sex differences in class II number in DBA, CAST, and 129S1 mice (male-female difference ± SE: 1.93 ± 0.58, z = 3.3; 1.75 ± 0.75, z = 2.33; and -2.94 ± 0.73, z = -4.01, respectively). Asterisks denote significant differences: **** p <0.0001, *** p<0.001, ** p<0.01, and * p<0.05 (Games-Howell for E, J; Bonferroni for L). Scale bars represent 10 µm. Error bars represent SD (E, J) or SEM (K).

Chiasmata counts were obtained from Giemsa-stained diakinesis/prometaphase I adult spermatocytes and juvenile oocytes (Fig. 1F-I, Fig. S1C-D, G-H, K-L, O-P. S-T).

For each sex and strain, the mean number of MLH1 foci per nucleus was quantified to estimate class I CO frequencies. One-way analysis of variance (ANOVA) indicated highly significant differences among groups (p < 0.0001; for full statistical comparisons, see Table S1). Females from most strains had significantly higher mean MLH1 foci counts by comparison to their respective males (Fig. 1E; Table 1). However, no significant sex difference in MLH1 foci was observed in 129S1 mice, while PWD mice exhibited an inverted sexual dimorphism with males averaging nearly five more MLH1 foci than females (p < 0.0001; Fig. 1E; Table 1, S1).

**Table 1.**
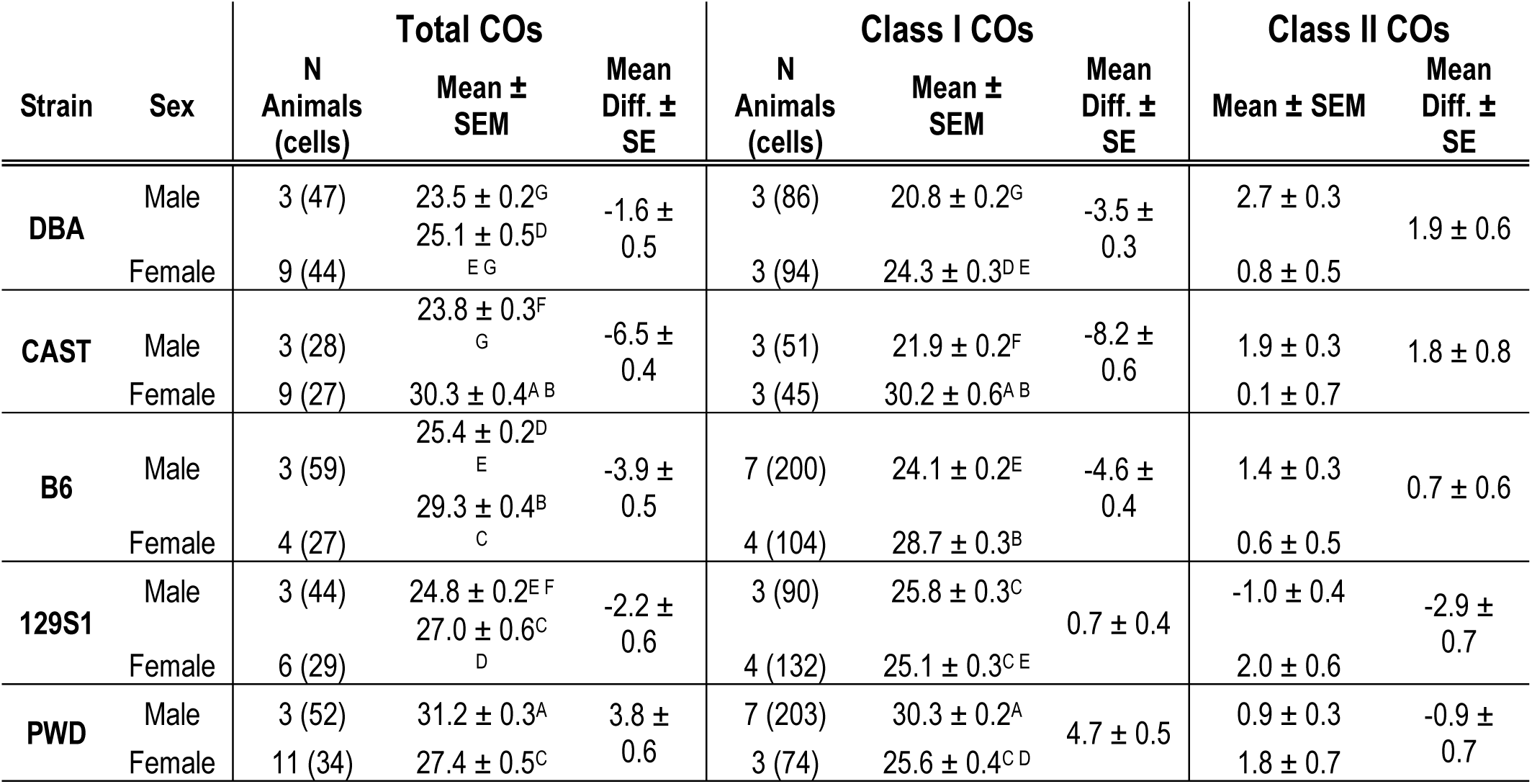
Crossover rates in male and female mice Mean total and class I crossovers quantified using per-nucleus counts of chiasmata and MLH1 foci, respectively. Class II crossover numbers were calculated by subtracting mean MLH1 foci from mean chiasmata. Superscript letters within a column denote significant differences as determined by one-way ANOVA and Games-Howell post hoc test (p<0.05); like letters indicate no difference.

Overall, PWD males had the highest mean MLH1 foci count of any sex and strain analyzed, while CAST mice showed the largest sex difference with females having 8.2 ± 0.6 more MLH1 foci than males (p < 0.0001, Fig. 1E, Table 1, S1). Chiasmata numbers generally corresponded to the strain-specific trends of sex-biased heterochiasmy we observed in MLH1 foci counts (p < 0.0001; Fig. 1J, Table 1; full statistical comparisons in Table S1). Notably, 129S1 females had more total chiasmata than their male counterparts (27.0 ± 0.59 and 24.82 ± 0.23, p < 0.05, Fig. 1J, Table 1, S1), making them the only strain to show a change in the directionality of heterochiasmy between male and female MLH1 and total CO counts. Prior studies have demonstrated that a minority (∼5-10%) of COs in mice are MLH1- independent and are presumed to be produced via the class II pathway, mediated by MUS81-EME1^34,62–65^. However, these studies were all conducted on the B6 background, and it is unknown whether proportions of class I and II COs are similar across mouse strains. Currently, a lack of commercially available antibodies viable for IF precludes direct cytological detection of class II COs. Thus, we estimated the frequency of class II COs by subtracting the mean number of MLH1 foci in pachynema from the total number of chiasmata (i.e., total CO number) and using the delta method to approximate their variance (Fig. 1K; Table 1). Within any given strain, the sex with the lower number of class I COs was estimated to have a higher relative number of class II COs (Fig. 1L). This trend held for all strains but only reached statistical significance for DBA (p < 0.001), CAST (p < 0.05), and 129S1 (p < 0.0001) mice (Fig. 1E, Table 1, S1). In the case of 129S1 males, the estimated number of class II COs was negative, highlighting that these values are estimates and should be interpreted with caution, and underscoring the differences in quantitation methods represented by immunofluorescence of surface spread chromosomes in prophase I and Giemsa staining of chromatin during diakinesis.

The low class I CO rates observed in CAST and DBA males (each approaching an average of 1 MLH1 focus per chromosome pair) raised the possibility that CO assurance may be more error-prone (i.e., less likely to make the obligate CO) in these mice. To test whether class II COs safeguard this obligate CO, we compared the percentage of pachytene cells containing at least one SC without an MLH1 focus (i.e., E0s, Fig. S2A) to the percentage of diakinesis/prometaphase I cells with at least one prematurely separated homologous pair (i.e., univalents; Fig. S2B). Chi-square analysis of the frequency of cells containing at least one E0 indicated highly significant differences among mouse strains (Χ^2^ = 69.0, p < 0.0001), and this occurrence was more common in females than males of the same strain (Fig. S2C). Cells with univalent chromosomes were not observed in either CAST or DBA males despite having two of the highest incidences of E0s among males. Notably, the only example of univalent chromosomes in males was observed in the 129S1 strain, which our estimates suggest make few or no class II COs (Fig. 1K, S2C). Taken together, these data suggest the class II pathway plays a critical role in ensuring that obligate COs are made on each chromosome pair and highlight the heterogeneous nature of class II pathway usage across divergent mouse strains.

### Sex differences in class I CO rates are not universally attributable to sex differences in SC length

Numerous studies using mouse models have shown compelling evidence to support two complementary hypotheses regarding the relationship between SC length and CO distribution:

(1) cells with longer SCs make more COs^45–47,50^ and (2) higher CO frequencies in eutherian females are attributable to longer female SC length^35,44,49,50^. It is thus logical to predict SCs to be significantly longer in PWD males compared females; similarly, 129S1 males and females would likely show no significant difference in SC length. To test these hypotheses, we measured total SC lengths (μm) by tracing SYCP3. Because the X and Y chromosomes exclusively pair and undergo reciprocal exchange at the pseudoautosomal region, they do not reflect the broader CO landscape ^66,67^. We therefore measured all SCs in oocytes but only autosomal SCs in spermatocytes. Although this creates a slight female bias in the analysis of total SC length, the inclusion of an “extra” SC in the female data is not sufficient to change our ultimate conclusions. To standardize our data to account for the “extra” female SC, we also measured the microns of SC length per MLH1 focus on a per-chromosome basis (Fig. 2B).

**Fig. 2.**
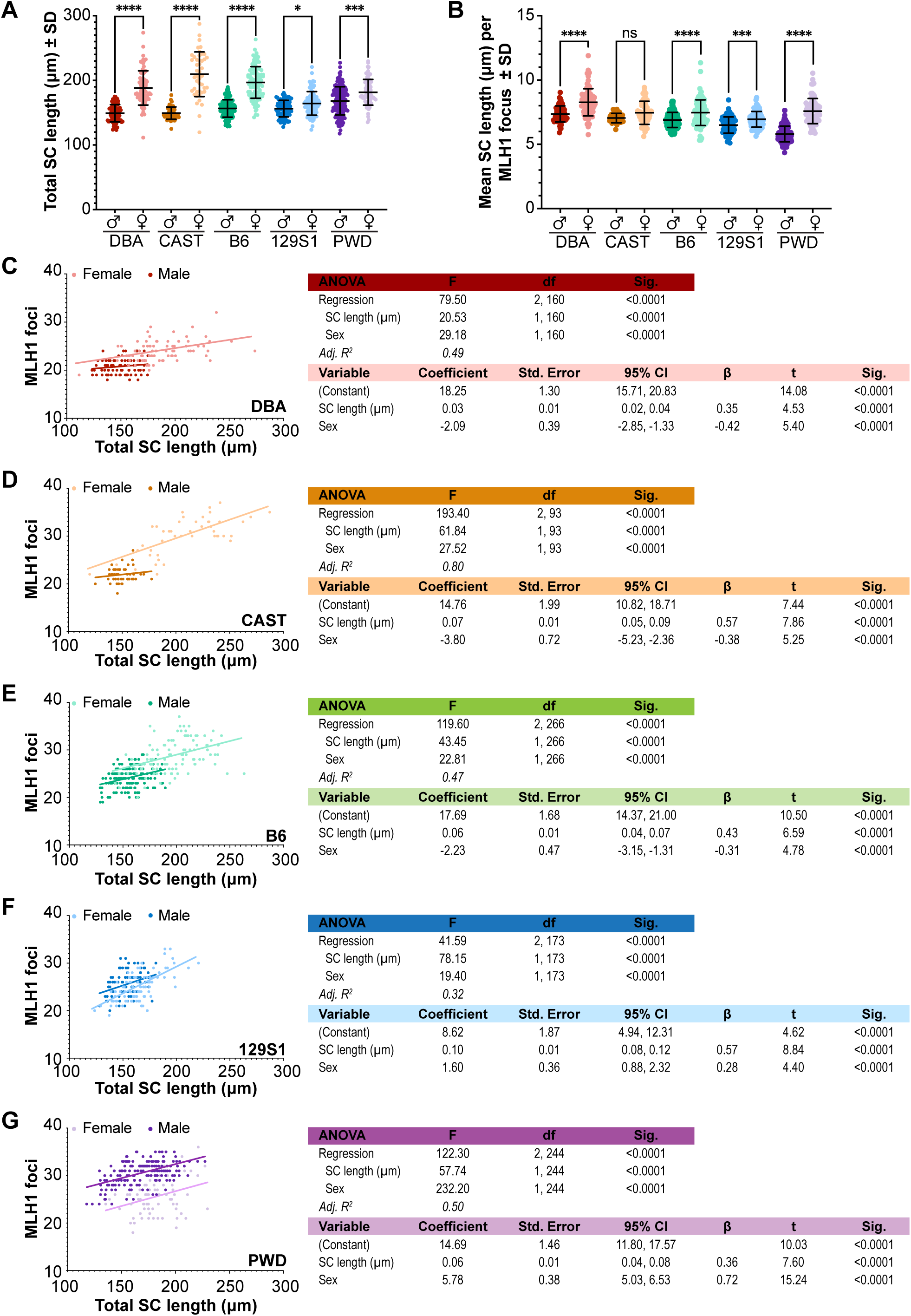
S**e**x **differences in SC length do not universally explain sex differences in crossover rate.** SC lengths and MLH1 foci position from the centromere were measured for a minimum of three animals per group and 20-30 cells per animal. (A) Females had significantly longer SCs than males in all mouse strains. (B) Except for CAST mice (no sex difference), males had fewer microns of SC length per MLH1 focus. (C-G) Multiple linear regression analysis of the effects of sex (male dark colors, female light colors) and total SC length on per- nucleus class I CO rates (MLH1 foci) for each strain: DBA, red (C); CAST, orange (D); B6, green (E); 129S1, blue (F); and PWD, purple (G). Adj. R^2^ denotes the proportion of variance in MLH1 foci number explained by SC length and sex (females as reference). Regression coefficients represent the change in MLH1 foci per one-unit change in either sex (male) or SC length (micron) holding other variables constant. Standardized regression coefficients (β) indicate the magnitude of each predictor’s effect (SC length or sex) on MLH1 foci number in common units (standard deviation). Asterisks denote significant differences: **** p <0.0001, *** p<0.001, and * p<0.05 (one-way ANOVA with Games-Howell post-hox; A, B). Error bars represent SD.

Comparing mean SC lengths indicated highly significant differences among sexes and strains (p < 0.0001; see Table S1 for full statistical analysis). For all strains, females consistently had significantly longer SCs than their respective males (Fig. 2A, Table 2, S1). Although the magnitude of the sex difference was less extreme in PWD mice than in other strains, it was still highly significant (p < 0.001; Fig. 2A, Table S1). Across strains, mice with the highest MLH1 focus counts also had the longest total SC length for their respective sex (e.g., the longest versus second longest SC lengths among males were PWD versus B6 mice, mean ± SEM: 168.3 µm ± 1.7 and 156.6 µm ± 1.1, respectively; p<0.0001; Fig. 2A, Table 2, S1). Our analysis of SC length per MLH1 focus indicated males had modestly, but significantly, fewer microns per focus by comparison to females (p<0.0001; Fig. 2B, Table 2; see Table S1 for full statistical comparison). Regardless of sex, most mice had between 7.0 and 7.5 μm /focus (Fig. 2B).

**Table 2.**
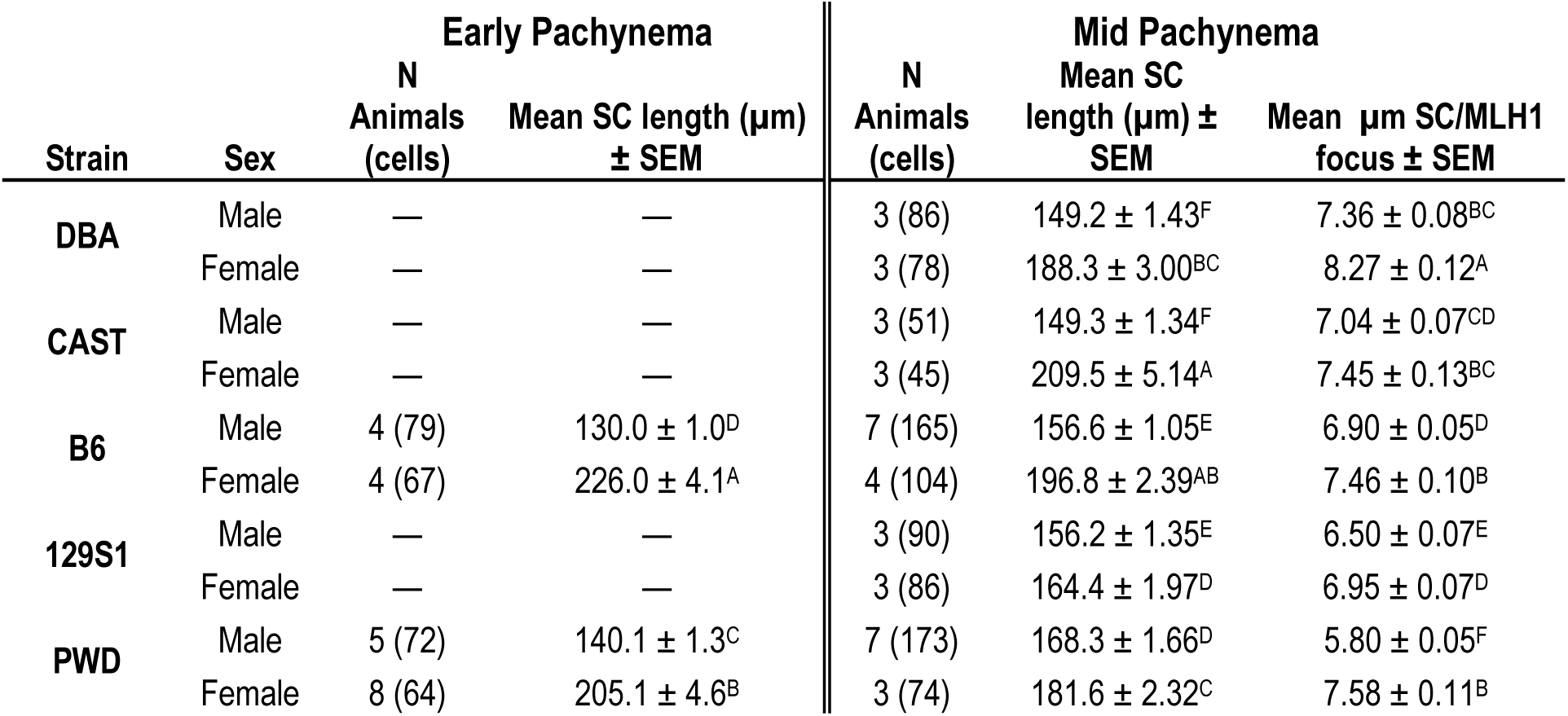
Total SC length per nucleus and microns of SC per MLH1 focus Mean SC length represents total length of all SCs per nucleus. Mean microns per MLH1 focus were calculated on a per-SC basis. Superscript letters within a column denote significant differences as determined by one-way ANOVA and Games-Howell post hoc test (p<0.05); like letters indicate no difference. — Denotes group analysis was not performed.

However, CAST females and PWD males were notable outliers (8.27 ± 0.12 μm /focus and 5.86 ± 0.05 μm /focus, respectively), with PWD mice showing the highest sex difference (males averaging 1.78 μm /focus less than females, (p < 0.0001; Table 2, S1).

To determine the relative effects of sex (female as the reference group) and SC length on MLH1 foci count, we performed a multivariable linear regression analysis (Fig. 2C-G). For all strains, the inclusion of a sex-SC length interaction did not significantly improve the model’s fit, indicating the effect of SC length on MLH1 foci was similar between males and females within each strain. The multiple regression was highly significant and explained ∼50% (i.e., Adj. R^2^ = 0.5) of the variation in MLH1 focus numbers for most strains (Fig. 2C, E, G). In contrast, the regression model showed very strong fit for CAST mice (explaining 80% of the variation in MLH1 focus numbers, Fig. 2D), but a more moderate level of fit for 129S1 mice (explaining 32% of variation, Fig. 2F). Thus, sex and SC length account for much of the variation in MLH1 focus numbers in most mouse strains, but they are less consequential (although still significant) factors in 129S1 mice.

For all strains, both SC length and sex were strong, highly significant predictors of MLH1 foci number; however, the magnitude and direction of their standardized effect sizes (β) varied considerably. Consistent with prior studies^46,47,50^, SC length universally had a positive effect on MLH1 foci number. The effect of sex on MLH1 focus numbers was also consistent with our observations of sex differences in crossover frequency (Fig. 1E), showing a negative effect of male sex among DBA, CAST, and B6 mice (Fig. 2C-E), but a positive effect for 129S1 and PWD mice (Fig. 2F, G). For CAST, B6, and 129S1 mice, SC length had a larger effect on MLH1 foci count than did sex; this difference was particularly striking in 129S1 mice, where β for SC length was 2-fold larger than β for sex (Fig. 2D-F, C, D). By contrast, for DBA and PWD mice, changes in MLH1 focus counts were attributable more to sex effects than SC length. For DBA mice this difference was subtle (1.2-fold increase over SC length, Fig. 2C), but the sex effect was much more pronounced in PWD mice (2-fold increase over SC length; Fig. 2G).

Although the interaction of sex and SC length was not predicted to significantly affect the regression model for any strain, sex and SC length did show moderate collinearity in DBA, CAST, and B6 mice (variance inflation factor [VIF] = 1.91, R^2^ = 0.48 for DBA mice; VIF = 2.52, R^2^ = 0.60 for CAST mice, and VIF = 2.13, R^2^ = 0.53 for B6 mice). In contrast, there was nearly no collinearity in 129S1 and PWD mice (VIF = 1.07, R^2^ = 0.06, VIF = 1.08, R^2^ = 0.08, respectively), suggesting that SC length and sex have nearly independent effects on MLH1 foci number in these strains. Taken together, this model suggests that, in contrast to strains like B6 and CAST, PWD mice do not exhibit parallel sexual dimorphism in SC length and class I COs, with males making more COs despite a shorter axis. Further, the assumption that sex mediates the effect of SC length on MLH1 foci number is not statistically supported in our analysis of PWD and 129S1 mouse strains.

### Males have stronger CO interference than females

A weaker CO interference resulting in narrower placement of adjacent MLH1 foci could provide an explanation for the exceptional sexual dimorphism in PWD mouse CO rates. To test this hypothesis, we analyzed three related parameters of interference: (1) Distribution of MLH1 foci along the length of the SC, (2) the distance between adjacent MLH1 foci, and (3) the evenness of MLH1 distribution.

### Distribution of MLH1 foci

To examine the sex- and strain-specific effects of interference on MLH1 distribution on SCs with two foci, we normalized measurements of MLH1 foci positions (as a fraction of SC length), beginning from the centromere. We then compared the likelihood that MLH1 foci were positioned in the terminal ends (i.e., < 0.25 or > 0.75) versus the central 50% of the SC (Fig. 3A). Across sexes and strains, the two MLH1 foci exhibited bimodal distribution, pushing placement of each focus towards opposing ends of the SC (Fig. 3D, G, J, M, P). Fisher’s exact test indicated that for all strains except 129S1 mice, males were significantly more likely than their corresponding females to place MLH1 foci in the terminal quartiles of the SC. Mice of the same sex were overall similar to each other across strains with two notable exceptions. Among females, PWD mice were the least likely to have terminal MLH1 foci, significantly less likely than B6 (p<0.001) and 129S1 (p<0.0001) mice. Among males, 129S1 mice were the least likely to have terminal foci, significantly less likely than PWD males (p<0.0001).

**Fig. 3.**
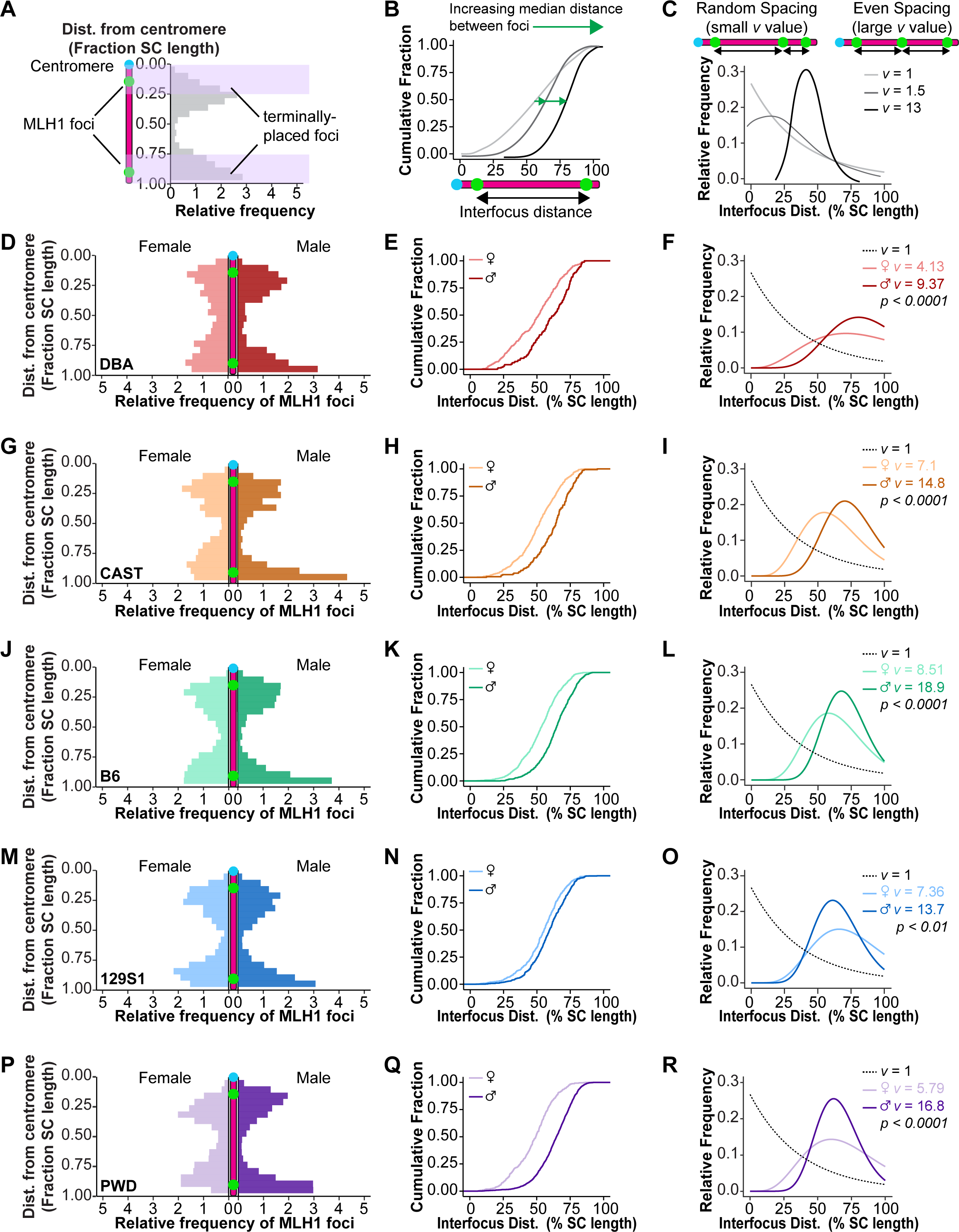
M**a**les **exhibit stronger crossover interference.** Three parameters were used to analyze CO interference strength: terminal placement of MLH1 foci (D, G, J, M, P), inter-focus distance standardized by %SC length (E, H, K, N, Q), and evenness of MLH1 foci spacing along the SC (F, I, L, O, R); interpretation overviews are shown in A-C, respectively. (A) The y-axis represents fractional SC length from centromere (0.0) to distal telomere (1.0); purple bars denote terminal SC ends (0.0–0.25 and 0.75–1.0). Histograms show relative frequency of MLH1 foci at each centromere distance. Males (dark bars) had significantly more terminally placed foci than females (light bars; Fisher’s exact test): 72.9% vs. 59.7% in DBA (D, OR = 0.55, p < 0.0001), 72.1% vs. 59.8% in CAST (G, OR = 0.58, p < 0.001), 72.9% vs. 61.8% in B6 (J, OR = 0.60, p < 0.0001), and 76.2% vs. 52.6% in PWD (P, OR = 0.35, p < 0.0001), but not in 129S1 (67.6% vs. 64.8%, OR = 0.88, p = 0.2). Among females, PWD mice had less terminally placed foci were than B6 (OR = 1.46, p<0.001) and 129S1 (OR = 1.66, p<0.0001). Among males, 129S1 mice had significantly fewer terminally placed foci than PWD (OR = 0.65, p<0.0001). (B) ECDF x-axis represents standardized inter-focus distance (%SC length); y-axis represents the fraction of SCs with inter-focus distance at or below the corresponding x-value. Steeper curves indicate tighter clustering; rightward shifts indicate greater median distances. Males had significantly greater median inter-focus distances than females (Kolmogorov-Smirnov with Bonferroni correction; Table S3) in DBA (E, KS = 0.26, p < 0.0001), CAST (H, KS = 0.31, p < 0.0001), B6 (K, KS = 0.34, p < 0.0001), 129S1 (N, KS = 0.13, p < 0.05), and PWD (Q, KS = 0.45, p < 0.0001). (C) Standardized inter-focus distances (%SC length) for each SC were fit to the gamma distribution using shape parameter *v*. Larger *v* values indicate more symmetrical, bell-shaped distributions and thus more regular MLH1 spacing (stronger interference); *v* = 1 (dotted line) represents an exponential distribution indicating random spacing and no interference. Males had significantly larger v than females in all strains: DBA (F) LR = 50.16; CAST (I) LR = 125.29; B6 (L) LR = 200.93; 129S1 (O) LR = 9.29; PWD (R) LR = 29.87.

To determine if chromosomes behaved differently based on their relative lengths, we also categorized SCs: the five longest (lower 25%, Fig. S4A) and five shortest (upper 25%, Fig. S4C). Long SCs showed the same trends found among pooled SCs (described above). Short SCs showed similar foci distributions regardless of genetic background, with no significant differences observed among males (Χ^2^ = 6.5, p = 0.16) or females (Χ^2^ = 7.1, p = 0.13) across strains. Females from most strains showed an increasing likelihood of terminal foci placement among short SCs relative to long SCs, while males showed similar foci distribution regardless of SC length (Fig. S4A, C). These trends erased sexual dimorphism of MLH1 distribution on short SCs in all strains except PWD mice (75.7% of foci were terminally placed in males vs. 55.6% in females, p<0.0001, Fig. S4C). However, short SCs with multiple MLH1 foci were too scarce in CAST and DBA males to reliably assess sex differences in those strains (Fig. S4C).

### Distance between MLH1 foci

To test whether the spacing between adjacent class I COs was greater in one sex, we examined the normalized distance (percentage of SC length) between MLH1 foci. The cumulative fraction of inter-focus distances was plotted (Fig. 3B) and significant differences between groups determined using Kolmogorov-Smirnov (KS) tests with a Bonferroni correction for multiple comparisons (Full statistical analysis in Table S3). For all mouse strains, males showed a significantly larger median spacing between adjacent MLH1 foci by comparison to females (Fig. 3E, H, K, N; Table S3). This difference was greatest in PWD mice (male versus female inter- focus distance of 64.7% vs. 49.3%, p<0.0001; Fig 3Q, Table S3). Across strains, males had similar inter-focus spacing except for 129S1 mice, which had significantly closer-spaced foci (59.40%) than either B6 (64.76%, p<0.0001)) or PWD males (p<0.0001, Table S3). Greater inter-strain variance was observed among females, with DBA (51.27%) and PWD (49.32%) mice having significantly narrower spacing by comparison to either 129S1 (55.39%; p<0.01) or B6 (53.50%; p<0.01) mice (Table S3). Consistent with our observations in MLH1 distribution, the longest five SCs per cell showed similar trends to the pooled analysis of inter-focus distances with males spacing foci significantly further apart than females (Fig. S4B). In contrast, MLH1 spacing on short SCs was similar across all sexes and strains except for PWD mice, where males again showed wider spacing between foci than females (63.2% vs. 47.0%, p<0.001; Fig. S4D, Table S3).

### Evenness of MLH1 distribution

The evenness of MLH1 spacing can be represented by the shape parameter (*v*) of distances between adjacent foci fitted to the Gamma-distribution, with larger values of *v* representing more symmetrical distributions and thus stronger interference (Fig. 3C). Because this analysis only accounts for chromosomes with multiple MLH1 foci, we employed statistical censoring to incorporate chromosomes with only one MLH1 focus in our analysis of interference. Measuring MLH1 distances from the centromere, the distance from the last (or only) focus to the end of the SC was considered right-censored while inter-focus distances were non-censored. Comparing the fitted Gamma-distributions between sexes for each strain using likelihood ratio tests indicated highly significant sex differences, with males exhibiting larger shape parameters (*v*) – and thus more even foci distributions – by comparison to females (Fig. 3F, I, L, O, R).

As most of our analyses of CO interference relied on SCs with at least two MLH1 foci, we wanted to determine if the frequency of SCs with multiple foci differed among sexes and strains. Of the five strains analyzed, only PWD males had multiple MLH1 foci on more than half their chromosomes (Fig. S3B). Short SCs with multiple class I COs were uncommon, accounting for less than 10% of short SCs for most groups. However, their frequency among PWD males was almost twice that of PWD females (16.5% vs. 8.4%) and nearly four times that of B6 males (4.4%; Fig. S3B). To estimate the likelihood of having multiple MLH1 foci on SCs at different lengths, we performed a logistic regression using SC length, sex, and strains, along with all two- and three-way interactions as predictors (Fig. S3A, C; Table S2). This model indicated that for DBA and CAST mice, females were significantly more likely to have multiple MLH1 foci on short (5µm) SCs (p < 0.0001 for both strains, Fig. S3A, C) while males were more likely on long (15 µm) SCs (p < 0.01 for DBA mice, p < 0.05 for CAST mice Fig. S3A, C).

Although B6 mice showed a similar trend, sex differences did not reach statistical significance at any SC length. Notably, only PWD males were significantly more likely than their corresponding females to have multiple MLH1 foci at all SC lengths (p < 0.0001; Fig. S3A, C).

Our data demonstrate that the evenness and spacing between MLH1 foci is greater in males than in females, leading chromosomes with multiple class I COs to have more terminally positioned COs in males than females. Taken together, this suggests that CO interference is stronger in males than in females. Consistently, PWD mice showed the most striking sex differences in MLH1 distribution and spacing. Although PWD males did trend towards having the highest likelihood of terminally positioned foci and the widest spacing between foci, this increased tendency was not significantly higher than in B6 males. By contrast, PWD females consistently exhibited more central MLH1 placement with narrower inter-focus spacing. Thus, the large sex difference in PWD MLH1 distribution was more likely an effect of females positioning a higher proportion of foci closer together and further from the SC ends. Thus, we reject our hypothesis that weaker interference in male mice explains the atypical sexual dimorphism of the PWD CO landscape. Instead, male PWD mice paradoxically have more class I COs spaced further apart on shorter axes by comparison to their female counterparts. Further, PWD males reliably make multiple class I COs, even on their shortest SCs, despite these limitations.

### Strong suppression of pericentromeric crossovers in PWD females

Pericentromeric suppression of crossing over is a well-documented phenomenon in meiosis ^5,68,69^ and is clearly observable in our MLH1 distribution data as a near absence of MLH1 across groups in the interval nearest the centromere (Fig. 3 D, G, J, M, P). To test whether pericentromeric crossover suppression differs among sexes and strains, we compared the normalized distance (%SC length) from the centromere to the first MLH1 focus (Fig. 4, full statistical analysis in Table S4). As described earlier, we were confined to exploring centromere effects on class I COs (MLH1 foci) because of the absence of reliable class II CO markers.

**Fig. 4.**
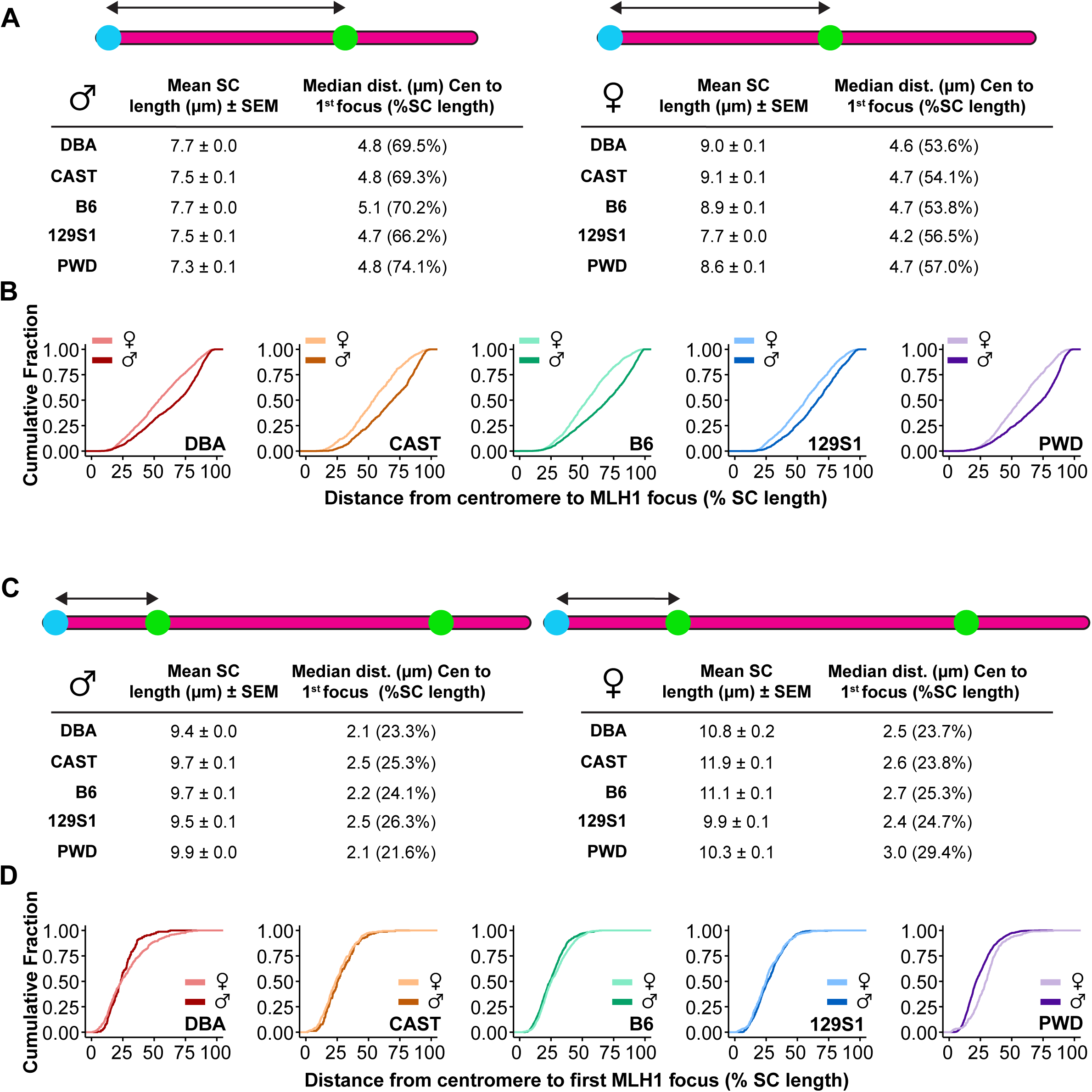
P**W**D **males exhibit weaker suppression of pericentromeric crossovers.** Pericentromeric CO suppression was measured as the normalized distance (%SC length) from the centromere to the first MLH1 focus for SCs with one (A-B) or multiple (C-D) MLH1 foci. (A, C) Diagrams illustrate the median distance (black double arrow) between the centromere (blue) and the first MLH1 focus (green) along the SC (magenta), with mean SC lengths shown. (B, D) ECDF x-axis represents standardized distance (%SC length) from centromere to first MLH1; y-axis represents the fraction of SCs with centromere-MLH1 distance at or below the corresponding x-value. Steeper curves indicate tighter clustering; rightward shifts indicate greater median distance. (B) For SCs with one MLH1 focus, males consistently had greater centromere distances than females in all strains (Kolmogorov-Smirnov: DBA, KS = 0.21, p < 0.0001; CAST, KS = 0.26, p < 0.0001; B6, KS = 0.27, p < 0.0001; 129S1, KS = 0.17, p < 0.0001; PWD, KS = 0.27, p < 0.0001). (D) For SCs with multiple MLH1 foci, significant sex differences in centromere-MLH1 distance were only observed in PWD (KS = 0.26, p < 0.0001).

Regardless of background, chromosomes with one MLH1 focus consistently exhibited significantly more telocentric MLH1 positioning in males by comparison to females (Fig. 4A-B). Across strains, PWD males and females tended to place their foci farther from the centromere than other strains (74.06% and 56.97% of the SC length, respectively), with PWD males showing significantly further placements than DBA (69.49%, p<0.0001) or 129S1 (66.22%, p<0.0001) males (Fig. 4A, Table S4). However, we reasoned that chromosomes with multiple MLH1 foci would best reveal the minimal distance of pericentromeric suppression because the centromere effect (pushing COs to the distal telomere) would be in opposition to interference (pushing COs toward chromosome ends). For chromosomes with multiple MLH1 foci, sex differences among strains largely vanished with the sole exception of PWD mice (21.60% vs. 29.43%, for males and females, respectively, p<0.0001; Fig. 4C-D; Table S4). Across strains, PWD males position their first MLH1 focus closest to the centromere, but the difference was only significant compared against 129S1 males (p<0.0001; Table S4). In contrast, the first MLH1 focus was spaced significantly farther from the centromere in PWD females than in all other strains (Table S4). These results indicate that the striking difference in centromere effect in PWD mice is primarily a result of stronger pericentromeric suppression of class I COs in PWD females.

### Sexually dimorphic dynamics of DSB repair differ between B6 and PWD mice

Having identified significant and unexpected sex differences in the PWD mouse CO landscape, we next examined the localization dynamics of proteins essential for DSB repair events leading to the formation of class I COs in order to ask whether specific steps in the pathway might show divergence between males and females. For the remainder of the studies described in this manuscript, we focused our analysis on PWD mice and use the well-characterized B6 mouse strain as a reference group.

Previous studies have suggested that strain differences in genome-wide CO rates are determined long before CO designation, with differences evident in the number of induced DSBs at the onset of meiosis^51,53^. To determine whether these observations held for males vs. females and between B6 and PWD mice, we used indirect IF staining against the RecA homolog, RAD51, which binds to the resected ends of all induced DSBs in zygonema to initialize homologous recombination (Fig. 5A-D)^70–72^. Analysis of mean counts of RAD51 foci indicated highly significant differences among groups (p<0.0001). Surprisingly, males of each strain did not significantly differ in RAD51 foci number (mean ± SEM: 213 ± 4.3 and 212.1 ± 4.4 for B6 and PWD, respectively; p = 1.0; Table 3, Fig. 5I). However, each strain exhibited opposing sexual dimorphisms: In B6 mice, females had considerably more RAD51 foci by comparison to males (306.6 ± 8.1, p<0.0001), while PWD females had fewer than their respective males (183.1 ± 5.9, p<0.001; Table 3, Fig. 5I). These data suggest that sex differences in crossing over for these strains might begin prior to or at the onset of DSB induction.

**Fig. 5.**
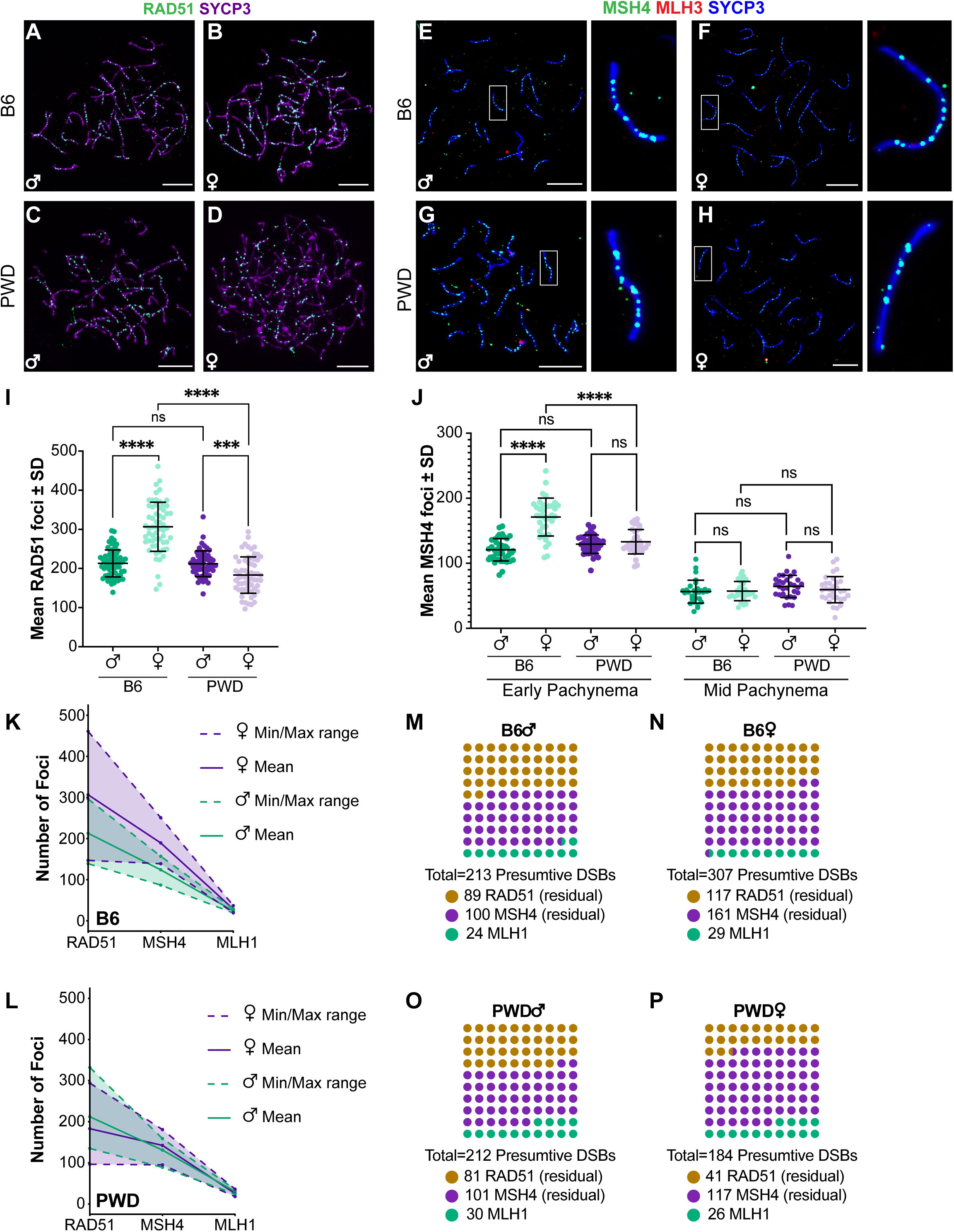
P**W**D **and B6 mice exhibit different sex-specific dynamics in the regulation of crossover designation.** Representative immunofluorescence images of zygotene spermatocytes and oocytes from B6 (A, B) and PWD (C, D) mice stained for RAD51 (green, proxy for presumed DSBs) and SYCP3 (magenta). Representative early pachytene spermatocytes and oocytes from B6 (E-F) and PWD (G-H) mice stained for MSH4 (green, proxy for CO intermediates), MLH3 (red, class I COs), and SYCP3 (blue). Enlarged views of SCs within white boxes are shown at right. (I) Comparison of mean RAD51 foci number (ANOVA F = 83.09, p<0.0001) indicates significant sex differences in B6 (t = 10.2) and PWD (t = 3.9). (J) Comparison of mean MSH4 foci number in early (ANOVA F = 43.71, p<0.0001) and mid-pachynema (F = 1.3, p = 0.3) indicated significant sex differences only in B6 at early pachynema (t = 9.12). (K-L) Progressive reduction from RAD51 (presumptive DSBs) to MSH4 (CO licensed sites) to MLH1 (designated class I COs) in male (green) and female (purple) B6 (K) and PWD (L) mice. Solid lines indicate mean foci number; dotted lines indicate minimum and maximum observed. (M-P) Presumptive DSB number (circles) and proportion of DSBs resolved as class I COs (green), converted to MSH4-marked licensed sites but not designated as COs (purple), or repaired by NCO pathways (orange) for B6 (M-N) and PWD (O-P) males and females, respectively. Asterisks denote significant differences: **** p < 0.0001, *** p < 0.001 (Games-Howell post-hoc; I, J). Scale bars represent 10 µm. Error bars represent SD.

**Table 3.**
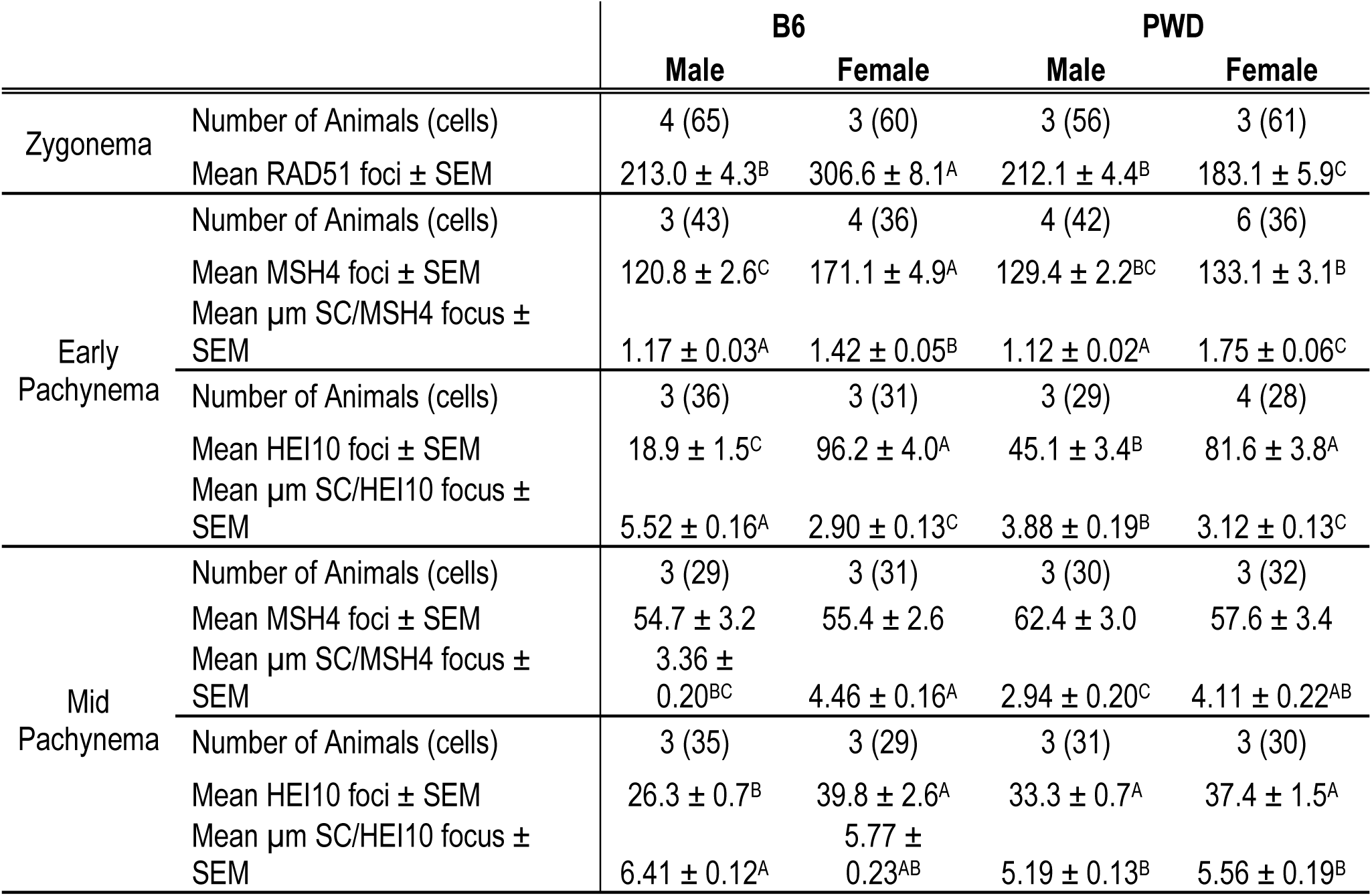
Mean foci number of RAD51, MSH4, and HEI10 foci throughout prophase I. Mean foci numbers were assessed per nucleus while mean microns per MSH4 or HEI10 focus were calculated on a per-SC basis. Superscript letters within a row denote significant differences as determined by one-way ANOVA and Games-Howell post hoc test (p<0.05); like letters indicate no difference.

|  |  | B6 |  | PWD |  |
| --- | --- | --- | --- | --- | --- |
|  |  | Male | Female | Male | Female |
| Zygonema | Number of Animals (cells) | 4 (65) | 3 (60) | 3 (56) | 3 (61) |
| | Mean RAD51 foci ± SEM | 213.0 ± 4.3<sup>B</sup> | 306.6 ± 8.1<sup>A</sup> | 212.1 ± 4.4<sup>B</sup> | 183.1 ± 5.9<sup>C</sup> |
| Early<br>Pachynema | Number of Animals (cells) | 3 (43) | 4 (36) | 4 (42) | 6 (36) |
| | Mean MSH4 foci ± SEM | 120.8 ± 2.6<sup>C</sup> | 171.1 ± 4.9<sup>A</sup> | 129.4 ± 2.2<sup>BC</sup> | 133.1 ± 3.1<sup>B</sup> |
| | Mean μm SC/MSH4 focus ± SEM | 1.17 ± 0.03<sup>A</sup> | 1.42 ± 0.05<sup>B</sup> | 1.12 ± 0.02<sup>A</sup> | 1.75 ± 0.06<sup>C</sup> |
|  | Number of Animals (cells) | 3 (36) | 3 (31) | 3 (29) | 4 (28) |
| | Mean HEI10 foci ± SEM | 18.9 ± 1.5<sup>C</sup> | 96.2 ± 4.0<sup>A</sup> | 45.1 ± 3.4<sup>B</sup> | 81.6 ± 3.8<sup>A</sup> |
| | Mean μm SC/HEI10 focus ± SEM | 5.52 ± 0.16<sup>A</sup> | 2.90 ± 0.13<sup>C</sup> | 3.88 ± 0.19<sup>B</sup> | 3.12 ± 0.13<sup>C</sup> |
| Mid<br>Pachynema | Number of Animals (cells) | 3 (29) | 3 (31) | 3 (30) | 3 (32) |
| | Mean MSH4 foci ± SEM | 54.7 ± 3.2 | 55.4 ± 2.6 | 62.4 ± 3.0 | 57.6 ± 3.4 |
| | Mean μm SC/MSH4 focus ± SEM | 3.36 ± 0.20<sup>BC</sup> | 4.46 ± 0.16<sup>A</sup> | 2.94 ± 0.20<sup>C</sup> | 4.11 ± 0.22<sup>AB</sup> |
|  | Number of Animals (cells) | 3 (35) | 3 (29) | 3 (31) | 3 (30) |
| | Mean HEI10 foci ± SEM | 26.3 ± 0.7<sup>B</sup> | 39.8 ± 2.6<sup>A</sup> | 33.3 ± 0.7<sup>A</sup> | 37.4 ± 1.5<sup>A</sup> |
| | Mean μm SC/HEI10 focus ± SEM | 6.41 ± 0.12<sup>A</sup> | 5.77 ± 0.23<sup>AB</sup> | 5.19 ± 0.13<sup>B</sup> | 5.56 ± 0.19<sup>B</sup> |

We next used indirect IF staining of the DNA mismatch repair protein MSH4 as a marker of CO licensing (Fig. 5E-H, Fig. S5A-D). Although peak MSH4 foci number has been reported to occur in late zygotene mouse spermatocytes^73^, we chose to analyze MSH4 in early pachynema in order to normalize counts by SC length in order to further examine the association between chromosome axis length differences and CO formation (Fig. S5E, Table 3). Comparison of mean total MSH4 foci number indicated highly significant differences (p<0.0001). In B6 mice, females had significantly more foci than males (mean ± SEM: 171.1 ± 4.7 and 120.8 ± 2.6, respectively, p<0.0001); however, in PWD mice there was no significant sex difference (129.4 ± 2.2 and 133.1 ± 3.1 for males and females, respectively, p = 0.9; Table 3, Fig. 5J). Sex differences in SC length were more pronounced in early pachynema than in mid, with females averaging 60 – 100 µm longer than males (Table 2, Fig. S6B). When MSH4 foci counts were weighted by SC length, clear sex differences in foci density were observed in both strains, with females consistently having more microns SC per focus by comparison to males (Table 3, Fig. S5E). By either metric (total MSH4 foci or weighted by SC length), males from each strain showed no significant difference.

We additionally analyzed MSH4 foci number in mid-pachynema (Fig. S5A-D, Table 3).

As pachynema progresses, MSH4 foci decline with an increasing proportion showing colocalization with class I CO markers while residual MSH4 is depleted as a consequence of non-CO DNA repair^30,73–75^. As expected, MSH4 foci were less numerous in mid-pachynema than at earlier stages, with all groups averaging from 54 – 63 ± 4 foci per nucleus. There was no significant difference between PWD and B6 males or females in total foci number (Fig. 5J, Table 3), but when weighting foci by SC length, males from each strain showed significantly higher densities of MSH4 than their corresponding females (Table 3, Fig. S5E). The lack of a significant difference between males of different backgrounds is suggestive of strain differences in the proportion of residual MSH4 foci in mid-pachynema. To test this, we examined the colocalization of MSH4 and the class I CO marker, MLH3, finding significant sex and strain differences. B6 females showed higher proportion of MSH4 foci colocalized with MLH3 than did their male counterparts (50.6% and 42.1%, respectively, p<0.0001; Fig. S5F), but PWD mice exhibited the inverse sexual dimorphism (males versus females: 47.9% and 41.7%, p<0.001; Fig. S5F).

Our analysis of mean RAD51, MSH4, and MLH1 foci numbers makes it possible to examine the overall trajectory of the class I CO pathway in males and females (Fig. 5K-L). To compare efficiency of establishing CO intermediates, we examined the ratios of MSH4 to RAD51 foci as a proxy for licensing efficiency using generalized linear mixed models (Fig. 5M- P). B6 males and females had similar ratios (0.57 and 0.56, respectively), but there was a much larger difference in PWD males and females (0.61 and 0.73). Comparison of the log-transformed ratios by generalized linear mixed modeling indicated that a significantly larger proportion of DSBs progress to CO intermediates in PWD females than in either PWD males (z = 3.02, p<0.05) or B6 females (z = -4.41, p<0.0001). No significant differences were observed among the other groups. Applying the same approach to MLH1:RAD51 as an estimate of the economical conversion of DSBs to mature class I COs, we found PWD mice had significantly larger proportions of DSBs resolve as class I COs for both males (0.14 and 0.11 in PWD and B6 mice, respectively; z = -5.76, p<0.0001) and females (0.14 and 0.09 in PWD and B6 mice, respectively; z = -8.05, p<0.0001). Within strains, only B6 mice showed a significant sex difference with males making more class I COs per DSB than females (z = -4.45, p<0.0001).

Finally, to estimate CO designation efficiency we compared MLH1:MSH4, finding highly significant sex and strain effects. Males of both strains consistently had larger ratios than their corresponding females (B6 males versus females: 0.20 and 0.17, z = -4.14, p<0.001; PWD males versus females: 0.23 and 0.19, z = -4.53, p<0.0001), and ratios in PWD mice were consistently larger than in B6 mice (males: z = -3.93, p<0.001; females: z = -2.66, p<0.05).

Taken together, these data suggest critical sex and strain differences in regulation and efficiency at multiple steps of crossing over: B6 and PWD mice exhibit opposite sexual dimorphism in DSB induction; sex differences in CO licensing efficiency are specific to PWD mice (females more efficient than males); but both strains show more efficient CO designation in males than in females.

### Sex- and strain-specific accumulation of the pro-crossover factor, HEI10, in pachynema

Finally, we wanted to identify a molecular explanation for the apparent sex and strain differences in class I CO maturation. Previous studies have identified several RING-E3 ligases, including HEI10, RNF212, and RNF212B, as critical to the stabilization of CO intermediates and designating which intermediates become class I COs^76–80^ – and these proteins have also been identified in QTL studies as contributing to inter-species differences in meiotic CO levels ^81–84^.

We thus hypothesized that differential RING-E3 ligase localization dynamics underpin sex and strain differences in crossover maturation. However, due to a lack of commercially available antibodies, we could only examine HEI10 localization by indirect IF (Fig. 6A-H).

**Fig. 6.**
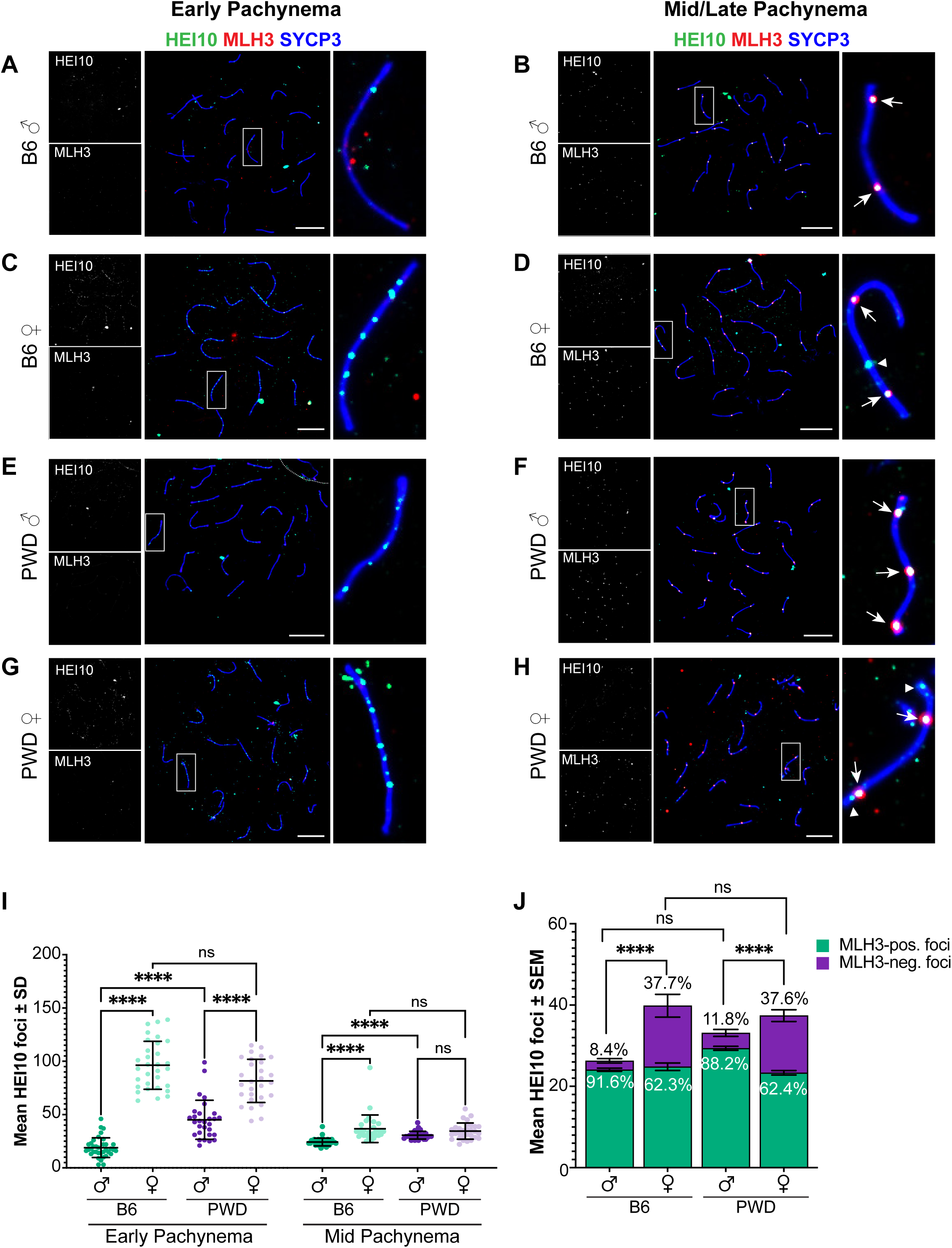
Sexual dimorphism in timing of HEI10 localization to the chromosome axis accumulation at presumptive class I crossover sites. Representative immunofluorescence images of early (A, C, E, G) and mid (B, D, F, H) pachytene spermatocytes and oocytes from B6 (A-D) and PWD (E-H) mice stained for HEI10 (green), MLH3 (red), and SYCP3 (blue). Individual HEI10 and MLH3 channels are shown in greyscale to the left of composite images; enlarged views of SCs within white boxes are shown at right. White arrows indicate HEI10 foci colocalized with MLH3; arrowheads indicate HEI10 foci without MLH3 colocalization. (I) Significant differences in HEI10 foci number at early pachynema (F = 118.7, p<0.0001) were observed between sexes in B6 (t = 17.88) and PWD (t = 7.12) mice; males also differed between strains (t = 7.00). At mid-pachynema (F = 15.14, p<0.0001), B6 males showed significantly fewer HEI10 foci than B6 females (t = 5.04) or PWD males (t = 7.17). (J) The proportion of HEI10 foci colocalized with MLH3 at mid-pachynema showed significant sex differences (B6: χ² = 234.8; PWD: χ² = 188.0) but no strain differences. Asterisks denote significant differences: **** p < 0.0001, *** p < 0.001 (Games-Howell for I; Bonferroni for J). Error bars represent SD (I) or SEM (J). Scale bars represent 10 µm.

In early pachynema we observed unexpected sex and strain differences in the timing of HEI10 localization to the SC. Females of each strain showed high numbers of HEI10 foci, markedly more than their ultimate number of MLH1 foci but fewer than early MSH4 (mean ± SEM: 96.2 ± 4.0 and 81.6 ± 3.8 for B6 and PWD females, respectively; Fig. 6I, Table 3). PWD males also exhibited a higher early HEI10 foci count than their eventual MLH1 foci count (45.1 ± 3.4), but significantly lower by comparison to PWD females (p<0.0001; Fig. 6I, Table 3). In contrast, B6 males had fewer HEI10 foci in early pachynema than their eventual mean MLH1 count (18.9 ± 1.5), and significantly fewer than either B6 females (p<0.0001) or PWD males (p<0.0001; Fig. 6I, Table 3). These trends were consistent whether analyzing total HEI10 foci per nucleus or weighting them by SC length (Fig. S6A, Table 3).

In mid-pachynema, HEI10 foci number increased in B6 males but decreased in all other groups. Previous studies have reported mid-pachytene spermatocyte HEI10 foci occur at near-identical levels as MLH1 in mice ^77,79,80^, and the males in our study were consistent with this. As such, significant differences in HEI10 foci number in mid-pachynema corresponded with differences in class I COs (Fig. 6I, Table 3). B6 males (26.3 ± 0.7) had significantly fewer HEI10 foci by comparison to B6 females (39.8 ± 2.6; p<0.001) and PWD males (33.3 ± 0.7, p<0.0001; Fig. 6I, Table 3). In contrast, B6 and PWD females (37.4 ± 1.5) did not significantly differ, and unexpectedly neither did PWD males and females (Fig. 6I, Table 3). As with our early pachynema HEI10 analysis, trends were consistent when foci counts were weighted by SC length (Fig. S6A). As females had more mid-pachynema HEI10 foci than COs, we examined the colocalization of HEI10 with a marker for class I COs, MLH3 (Fig. 6B, D, F, H, J). Males in both strains showed higher proportions of HEI10 colocalizing with MLH3 than did their female counterparts (B6 males versus females: 91.6% vs. 62.3%, p<0.0001; PWD males versus females: 88.2% vs. 62.4%, p<0.0001; Fig. 6J). Oocytes lack clearly defined fiduciary markers to reliably substage pachynema, thus there was a possibility the cells we analyzed for HEI10- MLH3 colocalization were not mid-pachytene or they were still in the process of CO designation. To rule out these possibilities, we compared MLH3 foci counts to our previous MLH1 foci counts and found them to be nearly identical (Fig. S6C), suggesting these cells had achieved the expected number of class I COs for their sex and strain. Thus, these findings indicate a relatively high number of residual HEI10 foci in oocytes that do not localize to sites of COs and lend support to our hypothesis of sex and strain differences in the regulation of CO designation.

## Discussion

Over the last twenty-five years, sexual dimorphism in SC length has been the leading explanation for heterochiasmy among mammals^35,44,49,50^. Our analysis of diverse mouse strains provides nuance to this long-standing dogma. We find that males from most backgrounds have shorter SC lengths and stronger CO interference than their corresponding females, and that these factors reinforce the lower rates of class I COs in these males relative to their female siblings. However, male PWD mice flout this rule and have higher CO frequency than females despite stronger interference and shorter axis length. Further, our analysis of key CO factors (RAD51, MSH4, and HEI10) indicates that sex and strain specific differences in the efficiency of CO designation underlie the exceptional sexual dimorphisms seen in PWD mice. Finally, our present study is the first to analyze both MLH1 foci in pachynema and chiasmata in diakinesis/metaphase I to evaluate sex differences in class I/II CO dynamics and distribution in mice from diverse genetic backgrounds.

### Interference and the centromere effect: Reinforcing the sexual dimorphic relationship between axis length and crossover rate

In meiosis, each chromosome is organized into a linear chromatin loop-axis array; sex differences in chromatin loop size impact the physical length of the axis. Interference further restricts the number of CO along the length of the axis. Although the mechanisms regulating CO interference remain poorly understood, it has been postulated that the designation of one CO at one location results in the propagation of an interference signal spreading in either direction away from the CO over a definite micron distance along the chromosome axis, thus CO number is thus predicted to positively scale with SC length^37,61,85,86^.

We predicted that the number of MLH1 foci would be limited by two complementary phenomena: (1) the centromere effect suppressing pericentromeric COs (pushing foci towards the distal end of the chromosome), and (2) CO interference causing wider spacing between adjacent MLH1 foci (pushing COs apart and toward the telomeres)^5,87,88^. Under this model, chromosomes with multiple MLH1 foci would require a sufficient axis length to accommodate both phenomena. Applying this framework to our data, we find that mean mouse SC length (7 – 10 µm) approximates the sum of the median inter-foci distance (5 – 6 µm) and pericentromeric suppression (2 – 3 µm), leaving little allowance for multiple MLH1 foci per chromosome. This likely explains why most mouse chromosomes across all strains and sexes – except PWD males – have only one MLH1 focus. Critically, we observed males across mouse strains exhibit stronger interference than their corresponding females. In context of their shorter chromosome axes, the greater inter-CO distance results in a lower likelihood of chromosomes having multiple MLH1 foci, thus greatly limiting CO frequency in males.

Pericentric suppression of crossing over is a second driving force of CO patterning. For single- MLH1 focus chromosomes, males show more distal CO placements than females across all strains, consistent previous reports^9,36,43,89–91^. For chromosomes with multiple MLH1 foci, however, sex differences in the centromere effect are absent in most strains. Notably, the centromere effect in PWD females was stronger than in either their corresponding males or females from other strains. Thus, the pronounced sex differences in CO positioning between PWD mice likely results from weaker interference combined with a stronger centromere effect in females. PWK/PhJ mice (close relatives of PWD/PhJ) possess larger minor satellite arrays compared to the other strains we analyzed^92,93^, and these ∼120 bp repeats define the core centromere where CENP-A binds^92,93^. Given strain relatedness, we predict PWD mice likely have enlarged minor satellite domains, potentially explaining the larger centromere effect.

However, why only PWD females show enhanced centromere effects remains unclear. Studies in *Drosophila* suggest centromeric repeat dosage does not alter the centromere effect^94^, rather pro-crossover factors mei-218 and rec and species-specific variants of the SC protein C(3)G have been shown to influence pericentric CO patterning^95–97^. Similarly, the cohesin subunit Psc3 in fission yeast and pericentromeric cohesin enrichment directed by phosphorylation of the Ctf19 kinetochore complex in budding yeast have been shown to suppresses crossing over^98–100^. Future investigation of sex-specific differences in pericentromeric chromatin architecture and cohesin distribution could reveal the molecular basis for sex and strain differences in the mouse centromere effect.

### What makes PWD mice so exceptional?

For most mouse strains, the data presented herein support the idea that CO interference and the centromere effect reinforce sex differences in SC length to drive heterochiasmy^35,44–47,49,50^. However, data presented here and elsewhere^53,54^ show that PWD mice clearly flout this dogma, with males exhibiting much higher CO number than females despite having shorter axes and stronger CO interference. Our regression modeling reveals an outsized and inverted effect of sex on MLH1 foci number specifically in PWD mice (i.e., males show large positive association with MLH1 number, whereas most other strains show smaller negative relationship). Moreover, sex and SC length have nearly independent effects on MLH1 focus numbers in PWD mice, demonstrating that factors beyond SC length sexual dimorphism determine heterochiasmy in this strain.

Using RAD51:MSH4 ratio as a proxy for CO licensing efficiency, we identified a sex difference in PWD but not B6 mice, indicating slightly higher licensing efficiency in PWD females. However, this difference only offset the initial sex difference in presumptive DSB induction. Rather, MSH4:MLH1 ratio analysis indicated higher CO designation is the primary driver of higher CO frequencies in PWD males versus females. Our data show sex differences in HIE10 localization dynamics in B6 mice consistent with recent reports^80^, while PWD males exhibit higher numbers of early HEI10 foci more similar to females. However, early HEI10 foci numbers do not correlate with strain-specific heterochiasmy patterns, as B6 and PWD females show similar HEI10 levels that exceed those seen in PWD males. Curiously, females from both strains show higher proportions of HEI10 foci in mid-pachynema that do not colocalize with class I COs (Fig 6D, H, J), which may reflect sexually dimorphic CO designation efficiency.

Our observations of CO dynamics in PWD mice reveal that chromosome axis length is neither a universal determinant of CO frequency, nor the only driver of heterochiasmy. HEI10 represents just one component in a complex regulatory network that varies by sex and genetic background and which might help to determine CO frequency and distribution along with many other key pro-CO factors that have been shown in B6 mice to exhibit sex differences in their regulation during prophase I (e.g. CNTD1, RNF212, and RNF212B)^78–80,101^. The lack of a causal relationship between chromosome axis length and heterochiasmy in PWD mice, therefore, highlights this strain as a critical model to further unravel the puzzle of sexual dimorphism in meiotic chromosome behavior.

### Sexually dimorphic balance between CO classes: Implications for CO homeostasis

Despite the very low rate of class II CO selection, this pathway appears to have a critical function in CO assurance as we observed unpaired univalents in diakinesis far less frequently than would be predicted from MLH1 focus counts through pachynema. This effect is particularly apparent in male mice of each strain. Rather than a universal sexual dimorphism in CO class usage rates, our data show that for a given strain, the sex with fewer class I COs tends to have more class II COs. These findings indicate that the two pathways are not independent and are likely regulated to ensure a balanced CO homeostasis. Previous studies using *Mus81-*null mice support this notion, finding that loss of class II COs elicits an upregulation of the class I pathway to maintain normal chiasmata number^34^. Our data also demonstrate that mice with higher class I rates — which also tend to have the longer SC lengths — have lower rates of class II COs.

Because class II COs are thought to be interference independent^6,34,102,103^, they are likely more restricted by the number of class I COs than by the axis length. This idea fits with recent work from our lab showing mice with defective MSH5 ATP binding lose all crossovers in diakinesis, suggesting that CO homeostasis likely involves the combination of both CO classes, with MutSγ serving as a hub for pathway selection^32^. Importantly, the high rate of MutSγ deposition in zygonema/early pachynema relative to the final MutLγ focus count in pachynema in mammals^30,32,73^ lends support to this conclusion, indicating that the selection of final CO sites, regardless of their classification, occurs from a supernumerary population of MutSγ-defined CO intermediates.

Taken together, these findings show that sexual dimorphism in crossing over results from the combined effects of structural and regulatory processes that vary across genetic backgrounds, indicating that no single mechanism can fully explain the diversity of CO outcomes in mice. Understanding how these mechanisms are coordinated in different sexes and genetic contexts will be critical for determining the evolutionary pressures that drive heterochiasmy and homeostasis between CO pathways.

## Materials and Methods

### Care and use of experimental animals

Breeding stocks of adult inbred C57BL/6J (000664), DBA/2J (000671), 129S1/SvImJ (002448), CAST/EiJ (000928), and PWD/PhJ (004660) mice were procured from The Jackson Laboratory. For each mouse strain, brother/sister breeding pairs were mated beginning at 6-weeks of age. Resulting offspring were weaned at 3-weeks of age, separated by sex, and housed in polysulfone cages (Allentown PC75JHT) on ventilated racks. No more than 5 mice were housed per cage. Each cage contained Bed-o’-cobs ¼” bedding (Scotts 330BB), Bio-Serv standard hut (Scotts K3352), and Crinkle Nest Kraft (Scotts CNK), and three 1” x 1” nestlets (Scotts NES7200) for enrichment. All mice were maintained in a climate-controlled environment with a 12-hr. light/dark cycle and provided food (Envigo Bioproducts Inc Rodent diet 7012) and water *ad libitum*. All adult and juvenile mice were euthanized using inhaled CO2 for at least 3 min. and monitored for cessation of breathing, followed by secondary internal cervical dislocation. All adult males in this study were euthanized between 10- and 15-weeks of age. Fetal mice were euthanized by decapitation. Juvenile female mice used for oocyte chiasmata analysis were euthanized between post-natal day (PND) 24 and 28. All fetal mice in this study were euthanized between 16.5- and 18.5-days post coitum (dpc) by decapitation. Animal handling and procedures were performed following approval by the Cornell Institutional Animal Care and Use Committee under protocol 2004–0063.

### Prophase I chromosome preparations

Meiotic chromosome preparations were made according to methods initially developed by Peters and colleagues and refined as described previously^104–106^. Testes were cleaned and detunicated in PBS. Seminiferous tubules were teased apart and incubated in hypotonic extraction buffer, pH 8.2-8.4 (30 mM Tris-HCl; 50 mM sucrose; 17 mM trisodium citrate dihydrate; and 5 mM EDTA) for 20 min. on ice. For every two slides made, approximately 7 mm of tubule was macerated in 60 µL of 500 mM sucrose at room temperature. Slides were coated in a thin layer paraformaldehyde (1% PFA, 0.15% Triton X, pH 9.2-9.3), and 20 µL of cell suspension was spread on each slide. Fetal ovaries were cleaned in PBS and incubated in hypotonic extraction buffer for 12-15 min. at room temperature. Both ovaries were macerated together in 35 µL of 500 mM sucrose. Slides were divided into three equivalent-sized square wells using a hydrophobic PAP pen and dipped into 1% PFA (0.15% Triton X, pH 9.2-9.3). For each square, 5 µL of cell suspension was spread across the section. Slides were incubated in a room temperature humid chamber for 2 hr. before being removed from the humid chamber and left to completely air-dry. Dried slides were washed for 2 min. in Milli-Q water with 0.4% Photo-flo 200 solution (Kodak Professional 1464510) and immediately immunostained.

### Immunofluorescence staining

Indirect immunofluorescence (IF) staining was performed as described previously^105,106^. Slides were washed for 10 minutes in PBS with 0.4% Photo-flo, then 10 minutes in PBS with 0.1% Triton X, and finally blocked for 10 minutes in 10% antibody dilution buffer (ADB; 100% stock consisted of 3% (w/v) bovine serum albumin [Millipore Sigma A5177-5EA], 10% (v/v) normal goat serum [Gibco 16210-072], 0.05% Triton X in PBS and sterilized using a 45 µm syringe filter). Primary antibodies were diluted in 100% ADB, applied to slides, and incubated in a room temperature humid chamber overnight. Slides were washed as before. Secondary antibodies were diluted and applied in a similar manner as the primary antibodies and incubated in a room temperature humid chamber for 2 hours in the dark. Slides were washed 3x 5 minutes in PBS with 0.4% Photo-flo, 1x 5 minutes in Milli-Q water with 0.4% Photo-flo and mounted using ProLong Diamond antifade with DAPI (Invitrogen P36962). Commercially available antibodies and dilutions used in this study are listed in Table S5, along with two custom antibodies previously made and used by the Cohen lab at Cornell University: guinea pig anti-MLH3^105^ and rabbit anti-SYCP3^27^.

### Chromosome preparations from diakinesis spermatocytes

Chromosome preparations from diakinesis spermatocytes were made according to methods developed by Evans and colleagues^107^ and as detailed previously^105^. Seminiferous tubules were placed in 2.2% sodium citrate and minced using scalpel blades for approximately 30 seconds. Tubules were transferred to a 15 mL conical tube and further dissociated by repeated pipetting. Samples were left to settle at room temperature for 15 minutes, and the supernatant was collected and centrifuged at 600xg for 5 minutes at room temperature. The cell pellet was resuspended in 4 mL of 1.1% sodium citrate and incubated at room temperature for 15 minutes. The sample was centrifuged as before, the supernatant discarded, and the pellet was vortexed into suspension as a fixative (75% methanol, 25% glacial acetic acid) was added dropwise. After two additional rounds of centrifugation and resuspension in the fixative, cells were dropped from approximately 2 m onto slides and air dried.

Oocyte diakinesis/prometaphase I chiasmata spreads were done as detailed previously^106^. Dictyate-arrested oocytes were isolated from the ovaries of unstimulated juvenile females. Ovaries were dissected into warmed, pre-gassed EmbryoMax M2 Medium (Millipore- Sigma MR-015) containing 2.5 µM milrinone (Millipore-Sigma M4659) to prevent meiotic resumption. Germinal vesicle (GV) oocytes were isolated using a 100 µm-wide Stripper Tip (Cooper Surgical MXL3-100) and collected in 20 µL drops of M2 medium with milrinone under EmbryoMax Filtered Light Mineral Oil (Millipore-Sigma ES-005). To induce meiotic resumption, denuded GV oocytes were washed by five serial passages through EmbryoMax Potassium Simplex Optimized Medium (KSOM; Millipore-Sigma MR-101-D). Washed GV oocytes were incubated in 20 µL KSOM drops under mineral oil (maximum 30 oocytes per 20 µL medium) at 37°C in 5% CO2 for 5 hrs. to reach diakinesis/prometaphase I. Oocytes were incubated in 0.9% sodium citrate hypotonic solution for 5 – 10 min before being transferred to a 1 µL drop of acidified water (8 drops glacial acetic acid in 50 mL ultrapure H2O) on a microscope slide. Excess liquid surrounding the oocyte was removed, and oocytes were burst by dropping fixative (75% methanol, 25% glacial acetic acid) from a 150 µm Stripper Tip (Cooper Surgical MXL3- 200) and air dried. All chiasmata spreads were stained with 4% Giemsa (Millipore-Sigma GS500), washed 3x 3 minutes in ddH2O, dried, and mounted with permount (Fisher Scientific SP15-100).

### Image acquisition

All slides were imaged using a Zeiss Axio Imager.Z2 with Colibri 7 epifluorescence microscope at 63x magnification. For immunofluorescence experiments, DAPI (385 nm), AF488 (475 nm), RR-X (555 nm), and AF647 (630 nm) were imaged sequentially using standardized exposure times for each antibody. Images were processed and adjusted using Zeiss Zen Blue version 3.0 (Carl Zeiss AG, Oberkochen, Germany) to standardize brightness and background across all images. SYCP3 and MLH3 were used to stage prophase I cells. In spermatocytes, the extent of X and Y chromosome synapsis was used to substage pachynema into early, mid and late, as described in previous work^108,109^. Pachytene oocytes from 17.5 dpc were considered early if all chromosome pairs were fully synapsed and MLH3 foci were present on 10 or fewer SCs. Slides stained with Giemsa were imaged using brightfield and processed using Zeiss Zen Blue version 3.0 (Carl Zeiss AG, Oberkochen, Germany).

### Synaptonemal complex measurement, foci quantification, and statistical analysis

For each chromosome pair (excluding the XY in spermatocytes), chromosome axis lengths were measured using Zeiss Zen Blue version 3.0 (Carl Zeiss AG, Oberkochen, Germany) by manually tracing SYCP3 beginning at the centromere (marked with CREST or DAPI-dense spots). For each cell, genome-wide SC length was calculated by summing lengths for all 20 chromosomes (for females) or all 19 autosomes (males). At least 20 cells were measured per animal and at least three animals per group (Table 2, S1). Among group differences in mean SC length were analyzed by Brown-Forsythe one-way ANOVA; statistically significant differences (p < 0.05) between groups were determined by Games-Howell’s multiple comparison test.

For MLH1, MSH4, HEI10, and RAD51, at least two independent observers who were blinded to the mouse strain manually counted foci using Photoshop (Adobe). Any focus that was at least 50% localized to SYCP3 was counted. MLH1, MLH3, MSH4, and HEI10 were additionally quantified by manually counting the fluorescence peaks for each antibody observed along tracings of chromosome axes (SYCP3). A flexible, manual thresholding of fluorescence intensity was used to define peaks as: (1) being clearly defined (at least 50% brighter than background signal), and (2) corresponding to visible foci as determined by manual comparison between images and fluorescence profile. Minor differences between foci and peak counts were reconciled on a per-focus basis, and cells with major scoring discrepancies, poor staining, or synaptic defects were excluded from analysis.

For MLH1, the distance from the centromere to each fluorescence peak was recorded for all SCs except the XY in spermatocytes. MLH1 foci were counted from 20 – 30 mid-pachytene cells per animal and at least 3 animals per group (Table 1-2, S1-4). Among group differences in mean MLH1 foci counts were analyzed by Brown-Forsythe one-way ANOVA; statistically significant differences (p < 0.05) between groups were determined by Games-Howell’s multiple comparison test. Correlations between SC length and MLH1 foci number was done for each sex and strain using simple linear regression. To assess the relationships between sex and SC length on MLH1 foci number, a multiple regression analysis was done for each mouse strain. For each regression analysis, significance of the model (p < 0.05) determined by F-test. To assess the distribution of MLH1 foci, SCs were grouped by whether they had 0, 1, or >1 foci. The proportion of SCs in each group were further categorized by length into “short,” “medium,” and “long” classes based on within-cell quartiles of total SC length (i.e., the longest and shortest five SCs and the intermediate-length 9-10 SCs). The relationship between SC length and MLH1 foci number were assessed by logistic regression performed in R (v4.5.1). Models were fitted using a binomial link function to assess the effects of sex, strain, and SC length (continuous or categorical) on the likelihood of multiple MLH1 foci per SC. Interaction terms were evaluated by likelihood ratio tests, and estimated marginal means were used to compare predicted probabilities among groups.

MSH4 and HEI10 foci in early and mid-pachynema were analyzed only for C57BL/6J and PWD/PhJ males and females. Colocalization of MSH4-MLH3 and HEI10-MLH3 in early and mid-pachynema was done by quantifying MSH4- or HEI10-exclusive, MLH3-exclusive, and colocalized foci/peaks. Foci were counted from 8 – 15 cells per substage per animal, and at least 3 animals per group (Table 3). Statistical analysis of both total foci counts and counts normalized by SC length were done for MSH4 and HEI10. For MSH4 data from each substage (exhibiting normal distribution), among group differences in mean foci counts were analyzed by Brown-Forsythe one-way ANOVA; statistically significant differences (p < 0.05) between groups were determined by Dunnett’s T3 multiple comparison test. For HEI10 data (not normally distributed); among group differences in median foci counts were analyzed by Kruskal-Wallis test; statistically significant differences (p < 0.05) between groups were determined by Dunnett’s multiple comparison test. Analysis of MHS4-MLH3 and HEI10-MLH3 co-localization was done by chi-square test with a Bonferroni correction for multiple comparisons.

Analysis of RAD51 foci was done only for C57BL/6J and PWD/PhJ males and females. RAD51 foci were quantified in zygotene cells with less than 50% synapsis. At least 3 animals were analyzed per group, with 15-20 cells per animal (Table 3). Among group differences in mean foci counts were analyzed by Brown-Forsythe one-way ANOVA; statistically significant differences (p < 0.05) between groups were determined by Games-Howell’s multiple comparison test.

Analyses of comparisons or each marker (RAD51 versus MLH1, MSH4 versus MLH1, and RAD51 versus MSH4) were performed in R (v4.5.1) using generalized linear mixed models (GLMM) with a Marker x Sex x Strain fixed-effects structure and a random intercept for mouse identity to account for within-animal correlation. GLMMs were initially fit with a Poisson link and assessed for overdispersion (Pearson Χ^2^/df) and those with a ratio > 1.2 were refit using a negative-binomial family. Estimated marginal means were obtained for each Marker x Sex x Strain combination and pairwise comparisons were assessed on the log scale with standard errors calculated using the delta method. Significant differences among group log-ratios were determined using z-tests with Bonferroni and false-discovery rate corrections for multiple comparisons.

### Quantification of chiasmata, calculation of class II crossover number, and statistical analysis

Quantification of chiasmata from all chromosome pairs (excluding the XY in spermatocytes) was done according to the previously published rubric^106^. Due to the limited yield of oocytes from juvenile ovaries, chiasmata were assessed from a minimum of 25 oocytes pooled from at least 3 individuals. For males, chiasmata were assessed from at least 3 individuals, 10-15 spermatocytes per individual. Among group differences in mean chiasmata counts were analyzed by Brown-Forsythe one-way ANOVA; statistically significant differences (p < 0.05) between groups were determined by Games-Howell’s multiple comparison test (Table 1, S1).

For each sex and strain, the number of class II crossovers was inferred by subtracting the mean MLH1 foci number from the mean chiasmata number for each group. The standard error (SE) was calculated by error propagation assuming independence between measures. Within each strain, male versus female differences in calculated class II crossover frequency were done by z-test based on the difference of means and pooled SE.

### Analysis of interference length, inter-MLH1 focus distances, and MLH1 focus to centromere length

For all SCs measured, the distance from the centromere to each MLH1 focus was recorded. An initial assessment of crossover interference was done by calculating the distances between MLH1 foci for SCs with more than one MLH1 focus. The cumulative fraction of inter-focus distances was measured both as a percentage of SC length and in microns. Differences in the probability distributions of inter-focus distances between strains and sexes were determined by Kolmogorov-Smirnov test, and statistically significant differences (p < 0.05) between groups were determined by Bonferroni post-hoc test (Table S3). To determine if inter-focus distance differed for long and short chromosomes, we repeated the above analyses using the five longest and the five shortest SCs per cell. To assess pericentromeric suppression of crossing over, this same statistical analysis was done using distance from the centromere to the first MLH1 focus – both for SCs with only one focus and for SCs with more than one focus (Table S4).

For SCs with two MLH1 foci, focus position distributions along the SC were analyzed in R (v4.5.1). For each SC, positions of individual MLH1 foci were normalized to SC length (0.0 – 1.0, beginning at the centromere), and the normalized positions were plotted as density-scaled histograms by sex and strain. Each focus was classified as “terminal” (≤ 0.25 or ≥ 0.75) or “interior” (0.25 – 0.75), and within-strain sex differences in the proportion of terminal versus interior foci was done using two-tailed Fisher’s exact tests; among strain differences were determined using chi-square tests with Bonferroni correction for multiple comparisons.

Overall crossover interference strength and the evenness of MLH1 spacing was calculated using R (v4.5.1) by fitting inter-focus distances to the gamma distribution and comparing shape parameter (*v*). Each SC contributed one right-censored interval (the distance from the last or only MLH1 focus to the distal end of the SC), while SCs with multiple MLH1 foci also contributed uncensored intervals (the distances between adjacent foci). Interval distances were normalized to SC length, and their distribution was fit to a gamma distribution using maximum likelihood estimation (MLE) under right-censoring, parameterized by shape (*v*) and scale (*θ*), as described previously^34^. Initial parameter estimates were obtained by method-of-moments and were optimized by Broyden–Fletcher–Goldfarb–Shanno (BFGS) minimization of the negative log-likelihood. Within each strain, likelihood ratio tests were used to compare sex-specific models (distinct *v*, shared *θ*) to pooled models (shared *v*, *θ*), and significance was assessed from the Χ^2^ distribution.

## Acknowledgements

We thank members of the Cohen lab for critical evaluation of this manuscript and for their input and insight throughout the course of these studies. We also extend special thanks to Mary Ann Handel for her invaluable feedback and advice during the preparation of this manuscript. We are grateful to the Cornell Center for Animal Resources and Education for providing animal care and veterinary services to research animals. Work described in this manuscript was supported by financial support from the Eunice Kennedy National Institute of Child Health and Development: K99112986 to TSH and awards HD041012 and HD097987 to PEC.

## Author contributions

Conceptualization: TSH. Investigation: TSH, AW, CPG. Formal Analysis: TSH, AW, ST. Resources: TSH, PEC. Writing – Original Draft Preparation: TSH. Writing – Review & Editing: TSH, PEC. Funding Acquisition: TSH, PEC

## Supporting Information

Fig. S1 Imaging of MLH1 and chiasmata Representative images of spermatocytes and oocytes for DBA (A-D), CAST (E-H), B6 (I-L), 129S1 (M-P), and PWD (Q-T) mice. (A-B, E-F, I-J, M-N, and Q-R) pachytene chromosome spreads stained for MLH1 (class I COs, green), SYCP3 (chromosome axis, magenta), and CREST (centromeres, blue). (C-D, G-H, K-L, O-P, S-T) Giemsa-stained chiasmata spreads from diakinesis/metaphase I chromosome spreads. Scale bars represent 10 µm.

Fig. S2 The class II pathway facilitates assurance of the obligate crossover. (A) Representative immunofluorescence image of a DBA oocyte in which one chromosome pair failed to make a class I CO (E0, white arrow). (B) Representative Giemsa-stained DBA oocyte chiasmata spread showing unpaired homologous chromosomes (univalents, white arrow). (C) Percentage of pachytene cells with at least one E0 SC (dark teal bars, left axis) and percentage of metaphase I cells with univalents (light teal circles, right axis). Although chromosome pairs without a class I CO—predicted to result in unpaired chromosomes at metaphase I that likely mis-segregate—are relatively common, the actual incidence of metaphase I cells with univalents is strikingly low across strains and sexes, indicating that in many meiotic cells, the obligate CO is made via the class II pathway. Scale bars, 10 µm.

Fig. S3 SC length affects sex differences in the likelihood of multiple crossovers per chromosome pair in most mouse strains. (A) Effect of SC length on the predicted probability of an SC having more than one MLH1 focus for DBA (red), CAST (orange), B6 (green), 129S1 (blue), and PWD (purple) males (dark) and females (light). Logistic regression analysis and significance are in Table S2. (B) Each SC per nucleus was classified as long (5 longest), short (5 shortest), or medium (9-10 intermediate-length). The percentage of SCs in each length category without an MLH1 focus (light teal), with 1 focus (teal), or with multiple foci (dark teal) is plotted for each sex and strain. (C) Logistic regression analysis of the effects of sex, strain, and SC length (continuous variable) on the likelihood of an SC having multiple MLH1 foci (B6 female as reference). This model predicted the likelihood of multiple MLH1 foci for SCs of three lengths: 5, 10, and 15 µm. Odds ratios below 1.0 indicate higher likelihood in males. Note that 15 µm SCs were exceptionally rare in males (absent in DBA males).

Fig. S4 CO interference is stronger in males from all strains for long SCs, but only in PWD males for short SCs. Expanded analysis from Fig. 3 examining CO interference on the longest (A-B) and shortest (C-D) five SCs per cell. Histograms (A, C) show the relative frequency of MLH1 foci at terminal SC ends (0.0–0.25 and 0.75–1.0). The y-axis represents fractional SC length from centromere (0.0) to distal telomere (1.0). (A) For long SCs, males (dark bars) had significantly more terminally placed MLH1 foci than females (light bars; Fisher’s exact test): 74.43% vs. 65.15% in DBA (OR = 0.64, p < 0.05), 69.44% vs. 58.05% in CAST (OR = 0.61, p < 0.05), 72.80% vs. 59.05% in B6 (OR = 0.54, p < 0.0001), and 74.61% vs. 55.60% in PWD (OR = 0.35, p < 0.0001), but not in 129S1 (66.92% vs. 63.14%, p = 0.3). (C) Among short SCs, only PWD males showed greater terminal placement (75.72% vs. 55.56%; OR = 0.40, p < 0.0001). No significant sex differences were observed in other strains (male vs. female: DBA, 56.25% vs. 52.17%, p = 0.79; CAST, 83.33% vs. 61.59%, p = 0.41; B6, 68.42% vs. 66.19%, p = 0.71; 129S1, 69.64% vs. 64.29%, p = 0.52). (B) ECDF graphs of standardized inter-focus distance (%SC length) for long SCs show significantly greater distances in males for all strains (Kolmogorov-Smirnov: DBA, KS = 0.32, p < 0.01; CAST, KS = 0.35, p < 0.0001; B6, KS = 0.40, p < 0.0001; 129S1, KS = 0.21, p < 0.01; PWD, KS = 0.43, p < 0.0001). (D) For short SCs, only PWD males showed significantly greater inter-focus distance than females (KS = 0.49, p < 0.001).

Fig. S5 Sex and strain differences in the density of MSH4 foci and their colocalization with MLH3 Representative images of mid-pachytene spermatocytes (A-B) and oocytes (C-D) from B6 (A, C) and PWD (B, D) mice stained for MSH4 (green), MLH3 (red), and SYCP3 (blue). (E) Comparison of mean microns of SC per MSH4 focus for B6 and PWD males and females at early (F = 45.11, p<0.0001) and mid-pachynema (F = 12.46, p<0.0001). Significant sex differences in mean microns/focus were observed in B6 (early: t = 4.14; mid: t = 4.29) and PWD (early: t = 9.91; mid: t = 3.98). Significant strain differences were only observed between females at early pachynema (t = 4.05). (F) The proportion of MSH4 foci colocalized with MLH3 at mid-pachynema showed significant sex differences (B6: χ² = 23.1; PWD: χ² = 13.9) and strain differences (males: χ² = 11.1; females: χ² = 27.5). Asterisks denote significant differences: **** p < 0.0001, *** p < 0.001, ** p < 0.01 (Games-Howell for E; Bonferroni for F). Error bars represent SD (E) or SEM (F). Scale bars represent 10 µm.

Fig. S6 Sex differences in HEI10 foci density, SC lengths differ between early and mid-pachynema, and MLH1 vs. MLH3 foci in mid-pachynema. (A) Mean microns of SC per HEI10 focus for B6 and PWD males and females at early and mid-pachynema. Significant sex differences in mean microns/focus were observed only at early pachynema in B6 (early: t = 12.6; mid: t = 2.5) and PWD (early: t = 3.25; mid: t = 1.64). Significant strain differences were only observed between males (early: t = 6.54; mid: t = 7.01). (B) Total SC length at pachynema (F = 134.3, p<0.0001) is shorter at early (closed circles) versus mid (open circles) stages for B6 (t = 10.09) and PWD (t = 11.42) males, but longer at early pachynema for B6 (t = 7.22) and PWD (t = 5.61) females. Total SC length is consistently greater in females than males for B6 (early: t = 22.66; mid: t = 11.02) and PWD (early: t = 12.71; mid: t = 3.42). (C) For B6 and PWD male and female mid-pachytene cells, mean MLH1 foci number (closed circles) does not differ significantly from mean MLH3 foci number (open circles). Asterisks denote significant differences: **** p < 0.0001, * p < 0.05 (Games-Howell post-hoc). Error bars represent SD.

Table S1 Full one-way ANOVA analyses Summary of Brown-Forsythe ANOVA test and subsequent Games-Howell’s multiple comparison test for significant differences in mean SC length, MLH1 foci number, microns SC per MLH1 focus, and chiasmata number between all sexes and strains.

Table S2 Logistic regression predicting the likelihood of an SC having multiple MLH1 foci B6 female mice were used as a reference group. Odds ratios greater than 1 indicate increased likelihood of an SC having more than one MLH1 focus with a change in the respective variable (increasing SC length or change in strain or sex) while holding all over variables constant. Model fits were evaluated using G² (likelihood ratio test statistic) and several pseudo-R² values. Both the main effects only model and the full model (with all interaction terms) showed significantly better fits than the null model (no predictors), and the full model showed marginally better fit than the main effects only model.

Table S3 Full analysis of distance between adjacent MLH1 foci Both the absolute (microns) and normalized (%SC length) distance between adjacent MLH1 foci on the same SC were analyzed by Kolmogorov-Smirnov (KS) test using pooled data from all SCs, the 5 longest SCs per cell, and the 5 shortest SCs per cell. Top table summarizes median inter-focus distance, and the bottom table summarizes the KS test statistics and their respective Bonferroni-adjusted p-values.

Table S4 Full analysis of distance from the centromere to the first MLH1 focus Both the absolute (microns) and normalized (%SC length) distance between the centromere and the first MLH1 focus for each SC were analyzed by Kolmogorov-Smirnov (KS) test using pooled data from all SCs. The analysis was also performed for SCs with 1 MLH1 focus and multiple MLH1 foci. Top table summarizes the median focus distance from the centromere, and the bottom table summarizes the KS test statistics and their respective Bonferroni-adjusted p- values.

Table S5 List of antibodies used in immunofluorescence staining experiments

