## Supplementary figures and images for "Chromosome length is not the sole determinant of sexually dimorphic crossover rates during mammalian meiosis: Insights from genetically diverse mouse strains"

### Fig. S1

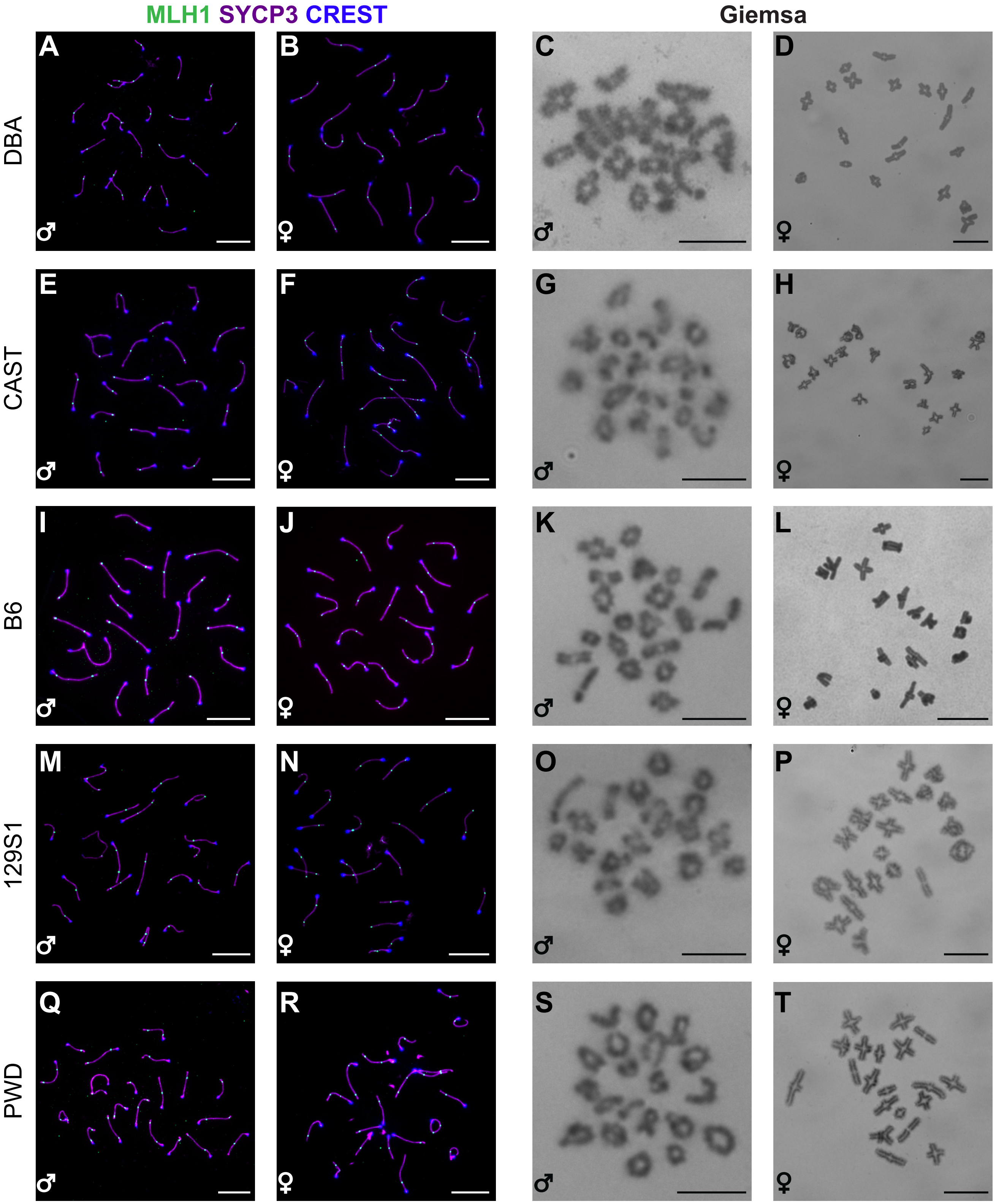

### Fig. S2

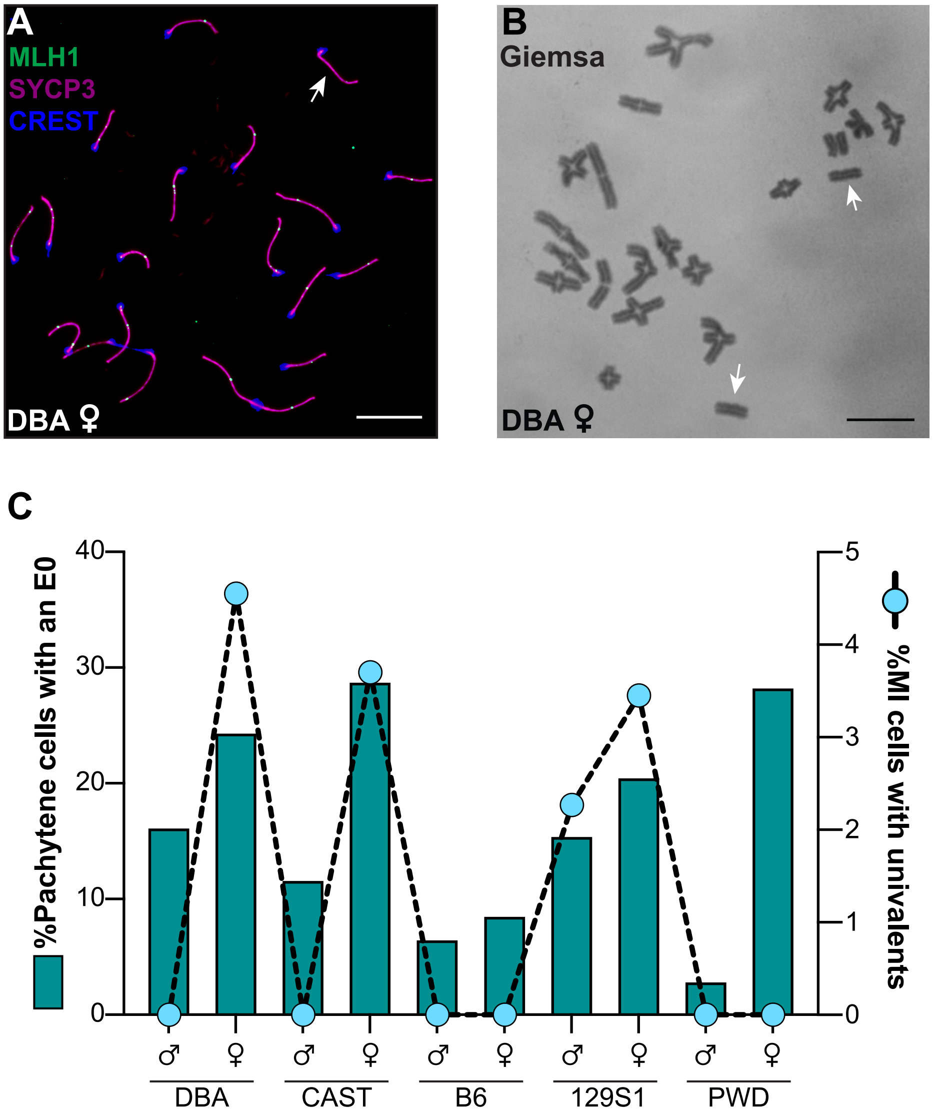

### Fig. S4

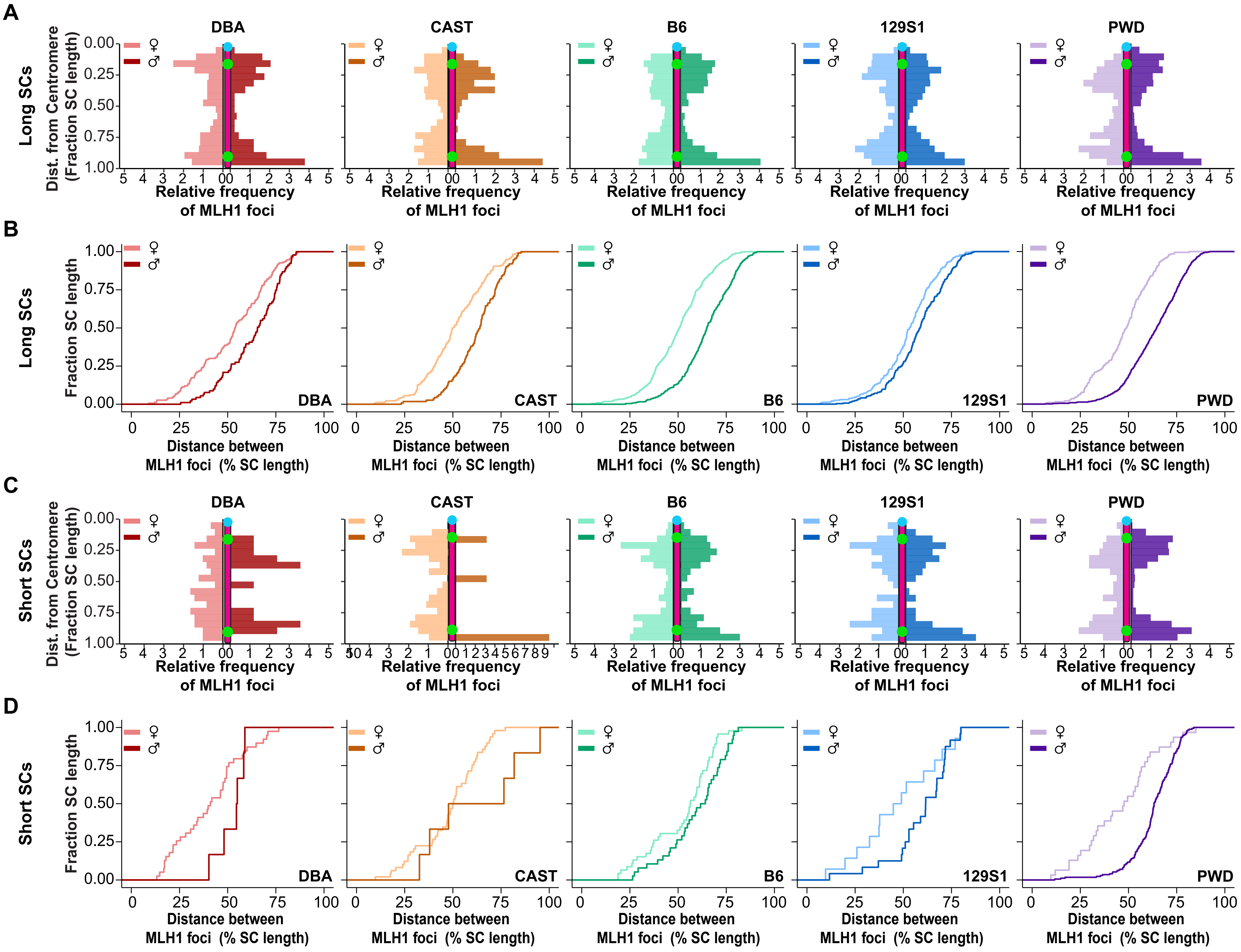

### Fig. S5

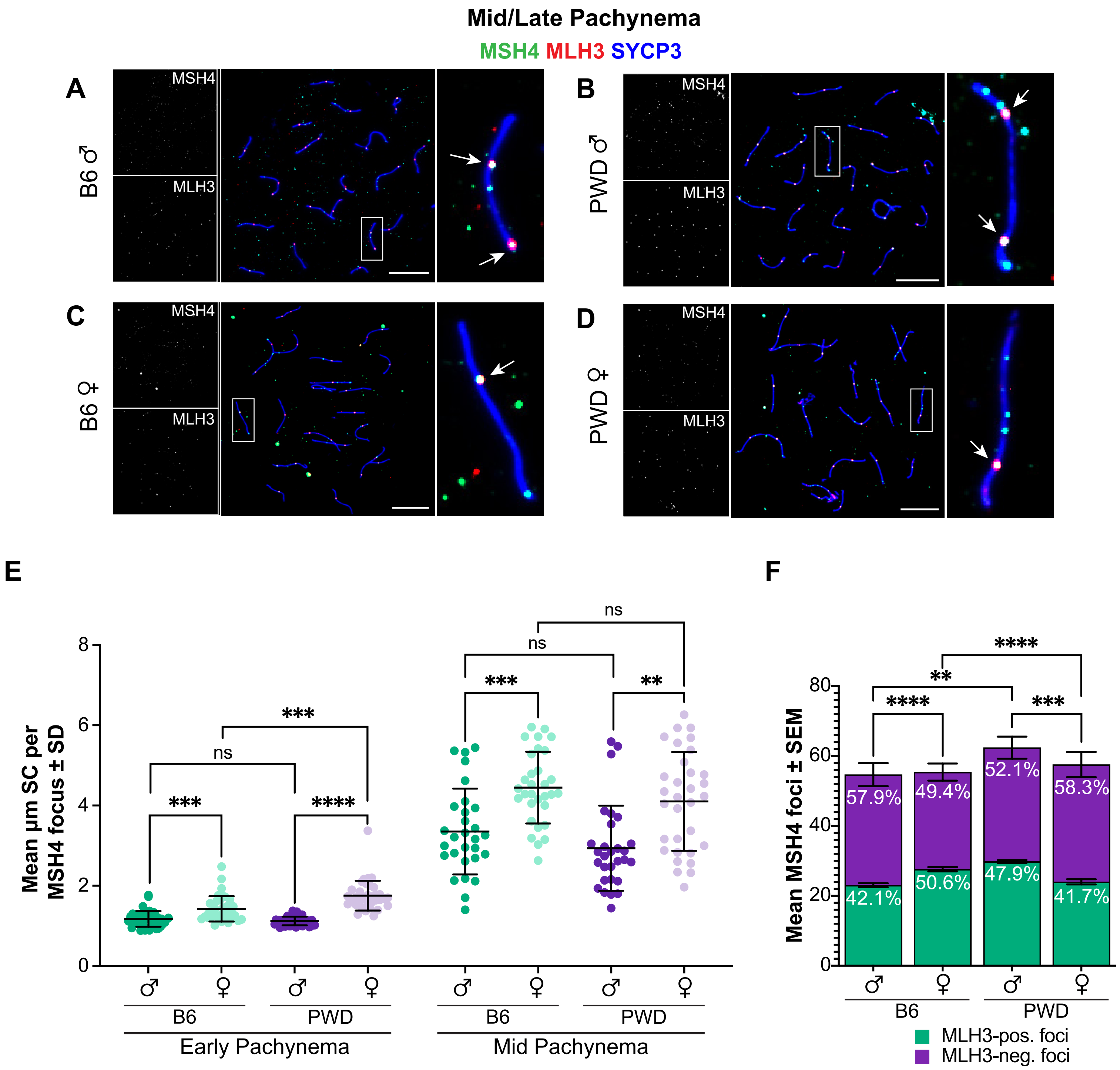

### Fig. S6

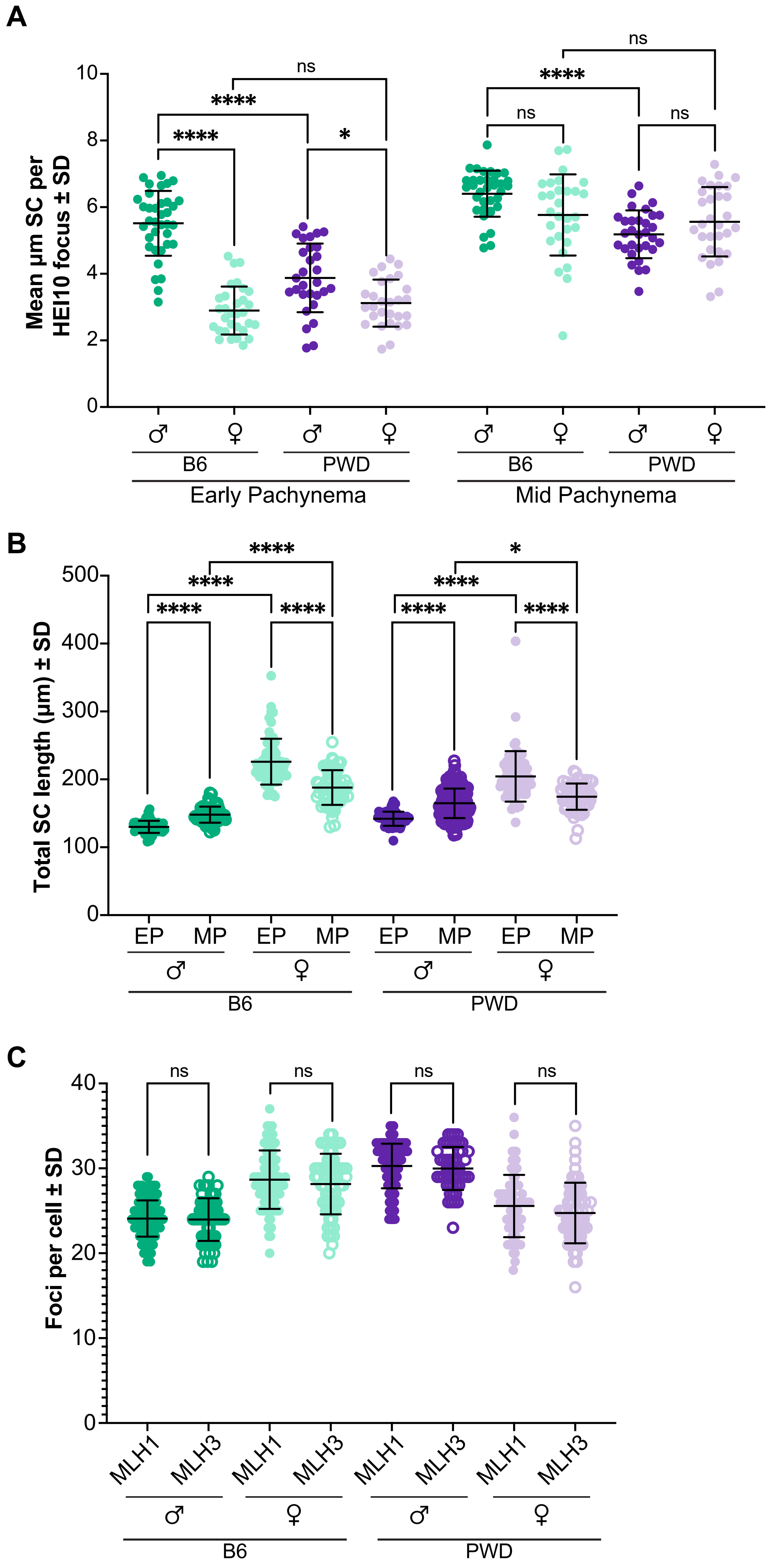
