## Supplementary material for "Chromosome length is not the sole determinant of sexually dimorphic crossover rates during mammalian meiosis: Insights from genetically diverse mouse strains": Fig. S3

A

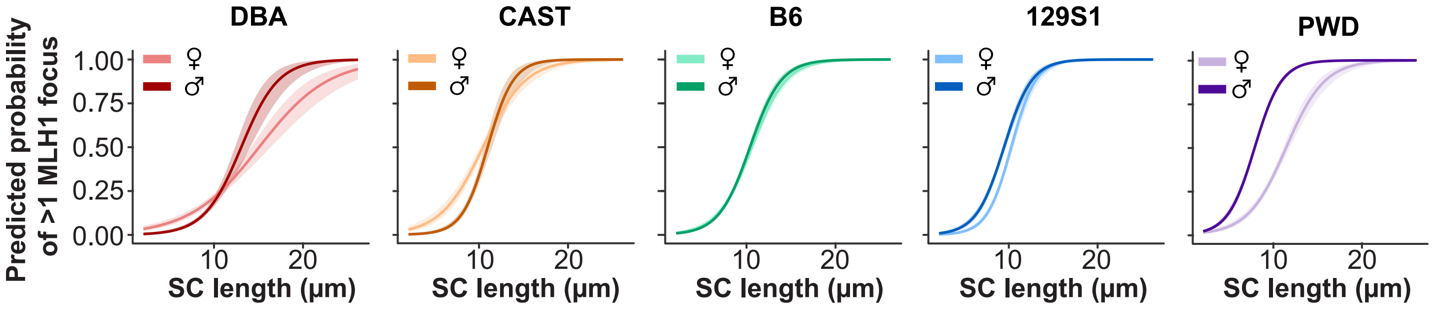


B

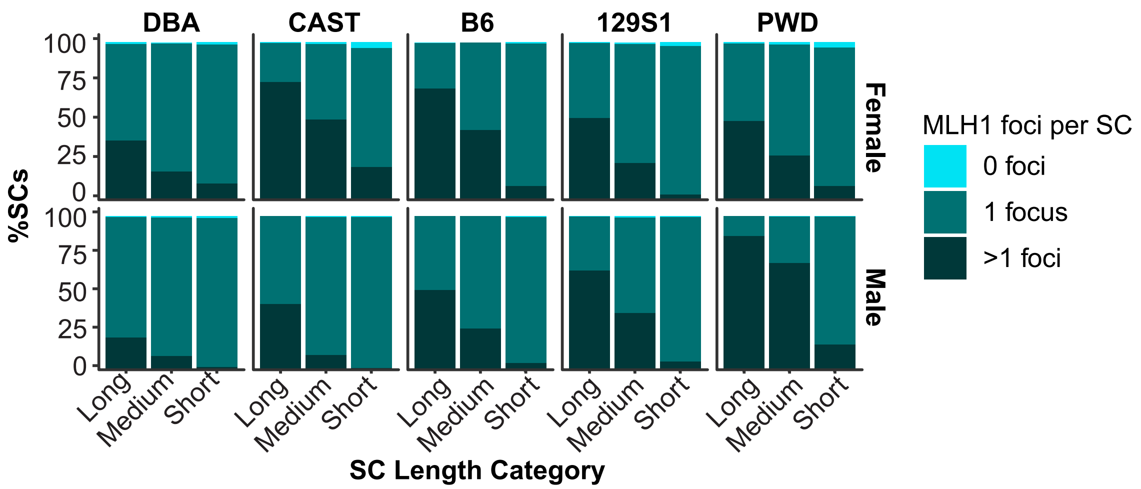


C

| SC Length (µm) | Strain | Odds Ratio | SE | z-ratio | p-value |
|---|---|---|---|---|---|
| 5 µm | DBA | 3.589 | 0.93 | 4.96 | <0.0001 |
| | CAST | 6.605 | 2.22 | 5.62 | <0.0001 |
| | B6 | 1.165 | 0.22 | 0.83 | 0.409 |
| | 129S1 | 0.381 | 0.09 | -4.16 | <0.0001 |
| | PWD | 0.362 | 0.07 | -5.12 | <0.0001 |
| 10 µm | DBA | 1.184 | 0.14 | 1.47 | 0.142 |
| | CAST | 1.670 | 0.23 | 3.78 | <0.001 |
| | B6 | 0.912 | 0.07 | -1.26 | 0.207 |
| | 129S1 | 0.583 | 0.06 | -5.01 | <0.0001 |
| | PWD | 0.129 | 0.01 | -23.39 | <0.0001 |
| 15 µm | DBA | 0.391 | 0.12 | -2.95 | <0.01 |
| | CAST | 0.422 | 0.18 | -2.07 | <0.05 |
| | B6 | 0.714 | 0.16 | -1.54 | 0.124 |
| | 129S1 | 0.892 | 0.31 | -0.33 | 0.745 |
| | PWD | 0.046 | 0.01 | -11.88 | <0.0001 |
