## Supplementary material for "Chromosome length is not the sole determinant of sexually dimorphic crossover rates during mammalian meiosis: Insights from genetically diverse mouse strains": Table S1

| | DBA/2J ♂ | DBA/2J ♀ | CAST/EiJ ♂ | CAST/EiJ ♀ | C57Bl/6J ♂ | C57Bl/6J ♀ | 129S1/SvImJ ♂ | 129S1/SvImJ ♀ | PWD/PhJ ♂ | PWD/PhJ ♀ |
|---|---|---|---|---|---|---|---|---|---|---|
| **N Animals (cells)** | 3 (86) | 3 (78) | 3 (51) | 3 (45) | 7 (165) | 4 (104) | 3 (90) | 3 (86) | 7 (173) | 3 (74) |
| **Mean SC length (µm) ± SEM** | 149.2 ± 1.43 | 188.3 ± 3.00 | 149.3 ± 1.34 | 209.5 ± 5.14 | 156.6 ± 1.05 | 196.8 ± 2.39 | 156.2 ± 1.35 | 164.4 ± 1.97 | 168.3 ± 1.66 | 181.6 ± 2.32 |
| **N Animals (cells)** | 3 (86) | 3 (94) | 3 (51) | 3 (45) | 7 (200) | 4 (104) | 3 (90) | 4 (132) | 7 (203) | 3 (74) |
| **Mean MLH1 foci ± SEM** | 20.78 ± 0.17 | 24.31 ± 0.26 | 21.94 ± 0.22 | 30.16 ± 0.57 | 24.09 ± 0.15 | 28.66 ± 0.34 | 25.77 ± 0.27 | 25.05 ± 0.26 | 30.28 ± 0.18 | 25.57 ± 0.43 |
| **N Animals (cells)** | 3 (86) | 3 (78) | 3 (51) | 3 (45) | 7 (165) | 4 (104) | 3 (90) | 3 (86) | 7 (173) | 3 (74) |
| **Mean microns SC per MLH1 focus ± SEM** | 7.37 ± 0.07 | 8.27 ± 0.12 | 7.08 ± 0.07 | 7.45 ± 0.13 | 6.91 ± 0.05 | 7.46 ± 0.10 | 6.50 ± 0.07 | 6.95 ± 0.07 | 5.86 ± 0.05 | 7.58 ± 0.11 |
| **N Animals (cells)** | 3 (47) | 9 (44) | 3 (28) | 9 (27) | 3 (59) | 4 (27) | 3 (44) | 6 (29) | 3 (52) | 11 (34) |
| **Mean chiasmata ± SEM** | 20.78 ± 0.17 | 24.31 ± 0.26 | 21.94 ± 0.22 | 30.16 ± 0.57 | 24.09 ± 0.15 | 28.66 ± 0.34 | 25.77 ± 0.27 | 25.05 ± 0.26 | 30.28 ± 0.18 | 25.57 ± 0.43 |

| ***Brown-Forsythe ANOVA test*** | ***SC length*** | ***MLH1 foci*** | ***Microns SC/MLH1 focus*** | ***Chiasmata*** |
|---|---|---|---|---|
| **F (DFn, DFd)** | 74.2 (9.0, 365.0) | 135.6 (9.0, 529.8) | 77.75 (9.0, 603.2) | 61.2 (9.0, 233.8) |
| **P value** | <0.0001 | < 0.0001 | <0.0001 | <0.0001 |

| ***Games-Howell's multiple comparisons test*** | ***SC length*** | | | | ***MLH1 foci*** | | | | ***Microns SC/MLH1 focus*** | | | | ***Chiasmata*** | | | |
|---|---|---|---|---|---|---|---|---|---|---|---|---|---|---|---|---|
| **Comparison** | **Mean Diff.** | **t** | **DF** | **Adj.P Value** | **Mean Diff.** | **t** | **DF** | **Adj.P Value** | **Mean Diff.** | **t** | **DF** | **Adj.P Value** | **Mean Diff.** | **t** | **DF** | **Adj.P Value** |
| DBA/2J ♂ vs. DBA/2J ♀ | -39.2 | 11.8 | 110.9 | <0.0001 | -3.5 | 11.6 | 157.4 | <0.0001 | -0.9 | 6.4 | 125.0 | <0.0001 | -1.6 | 3.2 | 59.0 | 0.0599 |
| DBA/2J ♂ vs. CAST/EiJ ♂ | -0.2 | 0.1 | 130.0 | >0.9999 | -1.2 | 4.2 | 104.6 | 0.0019 | 0.3 | 2.8 | 123.6 | 0.1350 | -0.3 | 0.9 | 56.7 | 0.9956 |
| DBA/2J ♂ vs. CAST/EiJ ♀ | -60.3 | 11.3 | 50.9 | <0.0001 | -9.4 | 15.7 | 51.6 | <0.0001 | -0.1 | 0.5 | 69.8 | >0.9999 | -6.8 | 16.7 | 42.8 | <0.0001 |
| DBA/2J ♂ vs. C57Bl/6J ♂ | -7.4 | 4.2 | 174.8 | 0.0018 | -3.3 | 14.7 | 218.6 | <0.0001 | 0.5 | 5.2 | 173.8 | <0.0001 | -2.0 | 6.6 | 104.0 | <0.0001 |
| DBA/2J ♂ vs. C57Bl/6J ♀ | -47.7 | 17.1 | 164.4 | <0.0001 | -7.9 | 20.9 | 148.6 | <0.0001 | -0.1 | 0.7 | 180.1 | 0.9996 | -5.8 | 12.8 | 38.8 | <0.0001 |
| DBA/2J ♂ vs. 129S1/SvImJ ♂ | -7.1 | 3.6 | 172.8 | 0.0155 | -5.0 | 15.8 | 148.2 | <0.0001 | 0.9 | 9.0 | 172.3 | <0.0001 | -1.3 | 4.4 | 86.8 | 0.0012 |
| DBA/2J ♂ vs. 129S1/SvImJ ♀ | -15.2 | 6.2 | 155.2 | <0.0001 | -4.3 | 13.9 | 207.1 | <0.0001 | 0.4 | 4.4 | 168.7 | 0.0008 | -3.5 | 5.7 | 34.6 | <0.0001 |
| DBA/2J ♂ vs. PWD/PhJ ♂ | -19.1 | 8.7 | 246.5 | <0.0001 | -9.5 | 38.3 | 257.9 | <0.0001 | 1.5 | 17.2 | 173.3 | <0.0001 | -7.7 | 23.2 | 92.3 | <0.0001 |
| DBA/2J ♂ vs. PWD/PhJ ♀ | -32.5 | 11.9 | 123.8 | <0.0001 | -4.8 | 10.5 | 95.1 | <0.0001 | -0.2 | 1.5 | 125.4 | 0.8714 | -3.9 | 7.1 | 43.0 | <0.0001 |
| DBA/2J ♀ vs. CAST/EiJ ♂ | 39.0 | 11.9 | 104.3 | <0.0001 | 2.4 | 7.0 | 140.0 | <0.0001 | 1.2 | 8.4 | 118.4 | <0.0001 | 1.3 | 2.5 | 64.5 | 0.2943 |
| DBA/2J ♀ vs. CAST/EiJ ♀ | -21.2 | 3.6 | 74.1 | 0.0218 | -5.8 | 9.3 | 62.3 | <0.0001 | 0.8 | 4.5 | 104.7 | 0.0006 | -5.2 | 9.0 | 69.0 | <0.0001 |
| DBA/2J ♀ vs. C57Bl/6J ♂ | 31.7 | 10.0 | 96.4 | <0.0001 | 0.2 | 0.7 | 159.6 | 0.9992 | 1.4 | 10.3 | 105.3 | <0.0001 | -0.3 | 0.7 | 62.9 | 0.9995 |
| DBA/2J ♀ vs. C57Bl/6J ♀ | -8.5 | 2.2 | 158.4 | 0.4460 | -4.4 | 10.3 | 187.1 | <0.0001 | 0.8 | 5.2 | 158.4 | <0.0001 | -4.2 | 6.9 | 67.8 | <0.0001 |
| DBA/2J ♀ vs. 129S1/SvImJ ♂ | 32.1 | 9.8 | 107.6 | <0.0001 | -1.5 | 3.9 | 181.3 | 0.0045 | 1.8 | 12.9 | 119.6 | <0.0001 | 0.3 | 0.5 | 62.9 | >0.9999 |
| DBA/2J ♀ vs. 129S1/SvImJ ♀ | 23.9 | 6.7 | 135.1 | <0.0001 | -0.7 | 2.0 | 218.4 | 0.5846 | 1.3 | 9.6 | 118.1 | <0.0001 | -1.9 | 2.6 | 58.3 | 0.2334 |
| DBA/2J ♀ vs. PWD/PhJ ♂ | 20.0 | 5.8 | 125.9 | <0.0001 | -6.0 | 18.9 | 190.6 | <0.0001 | 2.4 | 18.3 | 104.8 | <0.0001 | -6.1 | 11.6 | 70.2 | <0.0001 |
| DBA/2J ♀ vs. PWD/PhJ ♀ | 6.7 | 1.8 | 142.8 | 0.7529 | -1.3 | 2.5 | 122.7 | 0.2629 | 0.7 | 4.1 | 148.8 | 0.0023 | -2.3 | 3.4 | 71.6 | 0.0361 |
| CAST/EiJ ♂ vs. CAST/EiJ ♀ | -60.1 | 11.3 | 49.9 | <0.0001 | -8.2 | 13.4 | 56.6 | <0.0001 | -0.4 | 2.4 | 69.4 | 0.3376 | -6.5 | 14.8 | 48.2 | <0.0001 |
| CAST/EiJ ♂ vs. C57Bl/6J ♂ | -7.3 | 4.3 | 117.2 | 0.0015 | -2.1 | 8.1 | 103.5 | <0.0001 | 0.2 | 1.9 | 104.1 | 0.6758 | -1.7 | 4.9 | 64.6 | 0.0003 |
| CAST/EiJ ♂ vs. C57Bl/6J ♀ | -47.5 | 17.3 | 147.6 | <0.0001 | -6.7 | 16.7 | 152.2 | <0.0001 | -0.4 | 3.1 | 152.1 | 0.0760 | -5.5 | 11.5 | 44.4 | <0.0001 |
| CAST/EiJ ♂ vs. 129S1/SvImJ ♂ | -6.9 | 3.6 | 128.8 | 0.0146 | -3.8 | 11.1 | 138.1 | <0.0001 | 0.6 | 5.9 | 119.4 | <0.0001 | -1.0 | 3.0 | 61.4 | 0.1004 |
| CAST/EiJ ♂ vs. 129S1/SvImJ ♀ | -15.1 | 6.3 | 133.2 | <0.0001 | -3.1 | 9.2 | 165.4 | <0.0001 | 0.1 | 1.3 | 116.8 | 0.9406 | -3.2 | 5.1 | 38.5 | 0.0004 |
| CAST/EiJ ♂ vs. PWD/PhJ ♂ | -19.0 | 8.9 | 190.8 | <0.0001 | -8.3 | 29.2 | 130.8 | <0.0001 | 1.2 | 13.6 | 103.4 | <0.0001 | -7.4 | 20.0 | 71.6 | <0.0001 |
| CAST/EiJ ♂ vs. PWD/PhJ ♀ | -32.3 | 12.1 | 111.6 | <0.0001 | -3.6 | 7.6 | 105.5 | <0.0001 | -0.5 | 3.7 | 117.3 | 0.0124 | -3.6 | 6.3 | 48.2 | <0.0001 |
| CAST/EiJ ♀ vs. C57Bl/6J ♂ | 52.9 | 10.1 | 47.7 | <0.0001 | 6.1 | 10.3 | 50.3 | <0.0001 | 0.5 | 3.8 | 58.0 | 0.0136 | 4.8 | 11.6 | 47.1 | <0.0001 |
| CAST/EiJ ♀ vs. C57Bl/6J ♀ | 12.6 | 2.2 | 63.9 | 0.4496 | 1.5 | 2.2 | 76.1 | 0.4355 | 0.0 | 0.0 | 92.5 | >0.9999 | 1.0 | 1.8 | 51.0 | 0.7372 |
| CAST/EiJ ♀ vs. 129S1/SvImJ ♂ | 53.3 | 10.0 | 50.2 | <0.0001 | 4.4 | 7.0 | 63.8 | <0.0001 | 1.0 | 6.4 | 66.3 | <0.0001 | 5.4 | 13.0 | 47.0 | <0.0001 |
| CAST/EiJ ♀ vs. 129S1/SvImJ ♀ | 45.1 | 8.2 | 57.3 | <0.0001 | 5.1 | 8.1 | 63.0 | <0.0001 | 0.5 | 3.4 | 65.7 | 0.0393 | 3.2 | 4.7 | 45.5 | 0.0009 |
| CAST/EiJ ♀ vs. PWD/PhJ ♂ | 41.2 | 7.6 | 53.5 | <0.0001 | -0.1 | 0.2 | 53.5 | >0.9999 | 1.6 | 11.1 | 57.8 | <0.0001 | -0.9 | 2.1 | 54.8 | 0.5244 |
| CAST/EiJ ♀ vs. PWD/PhJ ♀ | 27.9 | 4.9 | 62.2 | 0.0003 | 4.6 | 6.4 | 89.8 | <0.0001 | -0.1 | 0.8 | 99.3 | 0.9991 | 2.8 | 4.6 | 55.8 | 0.0010 |
| C57Bl/6J ♂ vs. C57Bl/6J ♀ | -40.2 | 15.4 | 143.1 | <0.0001 | -4.6 | 12.4 | 145.2 | <0.0001 | -0.5 | 4.9 | 161.5 | <0.0001 | -3.9 | 8.3 | 42.0 | <0.0001 |
| C57Bl/6J ♂ vs. 129S1/SvImJ ♂ | 0.4 | 0.2 | 191.2 | >0.9999 | -1.7 | 5.5 | 147.9 | <0.0001 | 0.4 | 4.9 | 192.8 | <0.0001 | 0.6 | 2.0 | 98.0 | 0.6207 |
| C57Bl/6J ♂ vs. 129S1/SvImJ ♀ | -7.8 | 3.5 | 134.5 | 0.0227 | -1.0 | 3.2 | 218.2 | 0.0515 | 0.0 | 0.5 | 187.8 | >0.9999 | -1.6 | 2.5 | 36.1 | 0.2799 |
| C57Bl/6J ♂ vs. PWD/PhJ ♂ | -11.7 | 6.0 | 289.0 | <0.0001 | -6.2 | 26.0 | 386.9 | <0.0001 | 1.1 | 14.3 | 335.6 | <0.0001 | -5.8 | 16.7 | 102.8 | <0.0001 |
| C57Bl/6J ♂ vs. PWD/PhJ ♀ | -25.0 | 9.8 | 104.0 | <0.0001 | -1.5 | 3.3 | 91.9 | 0.0468 | -0.7 | 5.4 | 104.9 | <0.0001 | -2.0 | 3.5 | 45.5 | 0.0290 |
| C57Bl/6J ♀ vs. 129S1/SvImJ ♂ | 40.6 | 14.8 | 160.2 | <0.0001 | 2.9 | 6.7 | 187.4 | <0.0001 | 1.0 | 8.1 | 176.3 | <0.0001 | 4.5 | 9.6 | 42.2 | <0.0001 |
| C57Bl/6J ♀ vs. 129S1/SvImJ ♀ | 32.5 | 10.5 | 186.3 | <0.0001 | 3.6 | 8.5 | 204.4 | <0.0001 | 0.5 | 4.3 | 173.6 | 0.0011 | 2.3 | 3.2 | 49.1 | 0.0709 |
| C57Bl/6J ♀ vs. PWD/PhJ ♂ | 28.6 | 9.8 | 198.0 | <0.0001 | -1.6 | 4.2 | 166.0 | 0.0017 | 1.6 | 14.4 | 160.8 | <0.0001 | -1.9 | 3.9 | 48.4 | 0.0097 |
| C57Bl/6J ♀ vs. PWD/PhJ ♀ | 15.2 | 4.6 | 172.6 | 0.0004 | 3.1 | 5.7 | 151.1 | <0.0001 | -0.1 | 0.8 | 159.4 | 0.9981 | 1.9 | 2.9 | 58.3 | 0.1337 |
| 129S1/SvImJ ♂ vs. 129S1/SvImJ ♀ | -8.2 | 3.4 | 151.6 | 0.0272 | 0.7 | 1.9 | 209.3 | 0.6437 | -0.5 | 4.8 | 174.0 | 0.0001 | -2.2 | 3.5 | 36.4 | 0.0336 |
| 129S1/SvImJ ♂ vs. PWD/PhJ ♂ | -12.1 | 5.7 | 257.1 | <0.0001 | -4.5 | 13.9 | 176.2 | <0.0001 | 0.6 | 7.6 | 192.8 | <0.0001 | -6.4 | 18.3 | 93.5 | <0.0001 |
| 129S1/SvImJ ♂ vs. PWD/PhJ ♀ | -25.4 | 9.5 | 119.6 | <0.0001 | 0.2 | 0.4 | 125.7 | >0.9999 | -1.1 | 8.2 | 119.9 | <0.0001 | -2.6 | 4.6 | 45.8 | 0.0011 |
| 129S1/SvImJ ♀ vs. PWD/PhJ ♂ | -3.9 | 1.5 | 198.5 | 0.8832 | -5.2 | 16.5 | 254.7 | <0.0001 | 1.1 | 13.1 | 187.7 | <0.0001 | -4.2 | 6.5 | 39.7 | <0.0001 |
| 129S1/SvImJ ♀ vs. PWD/PhJ ♀ | -17.2 | 5.7 | 149.6 | <0.0001 | -0.5 | 1.0 | 127.2 | 0.9887 | -0.6 | 4.8 | 118.3 | 0.0002 | -0.4 | 0.5 | 58.3 | >0.9999 |
| PWD/PhJ ♂ vs. PWD/PhJ ♀ | -13.3 | 4.7 | 149.9 | 0.0003 | 4.7 | 10.2 | 101.5 | <0.0001 | -1.7 | 13.8 | 104.4 | <0.0001 | 3.8 | 6.6 | 50.7 | <0.0001 |
