## Supplementary material for "Chromosome length is not the sole determinant of sexually dimorphic crossover rates during mammalian meiosis: Insights from genetically diverse mouse strains": Table S2

| | Estimate | Std. Error | Odds Ratio | CI (low, high) | z-value | p-value |
|---|---|---|---|---|---|---|
| **(Intercept)** | -5.522 | 0.278 | 0.004 | 0.002 , 0.006 | -19.849 | <0.0001 |
| **SC Length (µm)** | 0.522 | 0.027 | 1.686 | 1.599 , 1.78 | 19.157 | <0.0001 |
| **Sex (Male)** | -0.397 | 0.367 | 0.672 | 0.327 , 1.384 | -1.082 | 0.2794 |
| **Strain (129)** | -1.843 | 0.476 | 0.158 | 0.061 , 0.398 | -3.875 | <0.001 |
| **Strain (CAST)** | 1.194 | 0.451 | 3.299 | 1.352 , 7.921 | 2.650 | <0.01 |
| **Strain (DBA)** | 1.650 | 0.386 | 5.209 | 2.443 , 11.112 | 4.273 | <0.0001 |
| **Strain (PWD)** | 0.301 | 0.442 | 1.352 | 0.565 , 3.199 | 0.682 | 0.4951 |
| **SC Length : Sex (Male)** | 0.049 | 0.038 | 1.050 | 0.974 , 1.13 | 1.292 | 0.1963 |
| **SC Length : Strain (129)** | 0.200 | 0.050 | 1.221 | 1.108 , 1.349 | 3.983 | <0.001 |
| **SC Length : Strain (CAST)** | -0.105 | 0.043 | 0.900 | 0.827 , 0.98 | -2.434 | <0.05 |
| **SC Length : Strain (DBA)** | -0.264 | 0.037 | 0.768 | 0.713 , 0.825 | -7.082 | <0.0001 |
| **SC Length : Strain (PWD)** | -0.065 | 0.044 | 0.937 | 0.86 , 1.022 | -1.462 | 0.1438 |
| **Sex (Male) : Strain (129)** | 1.788 | 0.621 | 5.977 | 1.774 , 20.258 | 2.879 | <0.01 |
| **Sex (Male) : Strain (CAST)** | -2.865 | 0.770 | 0.057 | 0.012 , 0.252 | -3.723 | <0.001 |
| **Sex (Male) : Strain (DBA)** | -1.989 | 0.629 | 0.137 | 0.039 , 0.464 | -3.163 | <0.01 |
| **Sex (Male) : Strain (PWD)** | 0.378 | 0.546 | 1.459 | 0.502 , 4.268 | 0.692 | 0.4888 |
| **SC Length : Sex (Male) : Strain (129)** | -0.134 | 0.067 | 0.875 | 0.766 , 0.997 | -1.993 | <0.05 |
| **SC Length : Sex (Male) : Strain (CAST)** | 0.226 | 0.080 | 1.254 | 1.073 , 1.47 | 2.819 | <0.01 |
| **SC Length : Sex (Male) : Strain (DBA)** | 0.173 | 0.065 | 1.189 | 1.046 , 1.351 | 2.650 | <0.01 |
| **SC Length : Sex (Male) : Strain (PWD)** | 0.158 | 0.057 | 1.171 | 1.046 , 1.309 | 2.769 | <0.01 |

| Model | log-likelihood | G2 | McFadden pseudo-$R^2$ | Cox & Snell's $R^2$ | Nagelkerke's $R^2$ | Chi-square | df | p-value |
|---|---|---|---|---|---|---|---|---|
| **Null** | -11770.567 | 0.000 | 0.000 | 0.000 | 0.000 | NA | NA | NA |
| **Main Effects** | -8767.115 | 6006.905 | 0.255 | 0.280 | 0.387 | 6006.90 | 15 | <0.0001 |
| **Full (with Interactions)** | -8752.043 | 6037.047 | 0.256 | 0.281 | 0.388 | 30.14 | 4 | <0.0001 |
