## Supplementary material for "Chromosome length is not the sole determinant of sexually dimorphic crossover rates during mammalian meiosis: Insights from genetically diverse mouse strains": Table S3

| | Measurement | Median Interfocus Distance DBA/2J ♂ | DBA/2J ♀ | CAST/EiJ ♂ | CAST/EiJ ♀ | C57Bl/6J ♂ | C57Bl/6J ♀ | 129S1/SvImJ ♂ | 129S1/SvImJ ♀ | PWD/PhJ ♂ | PWD/PhJ ♀ |
|---|---|---|---|---|---|---|---|---|---|---|---|
| All SCs | Absolute distance (µm) | 5.65 | 5.43 | 6.15 | 5.92 | 6.09 | 5.79 | 5.52 | 5.41 | 6.14 | 4.98 |
| | Normalized distance (%SC length) | 63.00 | 51.27 | 64.27 | 51.91 | 64.76 | 53.50 | 59.40 | 55.39 | 64.74 | 49.32 |
| Longest 5 SCs | Absolute distance (µm) | 6.84 | 6.67 | 6.43 | 6.80 | 7.02 | 6.28 | 6.43 | 5.85 | 7.23 | 5.48 |
| | Normalized distance (%SC length) | 65.88 | 53.82 | 64.03 | 49.99 | 64.62 | 50.87 | 59.22 | 53.18 | 64.91 | 48.94 |
| Shortest 5 SCs | Absolute distance (µm) | 3.55 | 2.89 | 2.54 | 4.22 | 4.33 | 4.38 | 4.42 | 3.60 | 4.76 | 3.26 |
| | Normalized distance (%SC length) | 54.98 | 41.07 | 47.73 | 50.05 | 64.53 | 56.34 | 67.21 | 46.91 | 63.17 | 46.98 |

| | All SCs | | | | Longest 5 SCs | | | | Shortest 5 SCs | | | |
|---|---|---|---|---|---|---|---|---|---|---|---|---|
| | Absolute Distance (µm) | | Normalized Distance (%SC Length) | | Absolute Distance (µm) | | Normalized Distance (%SC Length) | | Absolute Distance (µm) | | Normalized Distance (%SC Length) | |
| Comaprison | KS Stat | Adj. P value | KS Stat | Adj.P value | KS Stat | Adj. P value | KS Stat | Adj.P value | KS Stat | Adj. P value | KS Stat | Adj.P value |
| DBA ♀ vs. DBA ♂ | 0.13 | 1.00 | 0.26 | <0.0001 | 0.14 | 1.00 | 0.32 | >0.01 | 0.38 | 1.00 | 0.59 | 1.00 |
| CAST ♀ vs. CAST ♂ | 0.12 | 1.00 | 0.31 | <0.0001 | 0.22 | 0.12 | 0.35 | >0.0001 | 0.40 | 1.00 | 0.38 | 1.00 |
| B6 ♀ vs. B6 ♂ | 0.09 | <0.05 | 0.34 | <0.0001 | 0.20 | <0.0001 | 0.40 | >0.0001 | 0.14 | 1.00 | 0.27 | 1.00 |
| 129 ♀ vs. 129 ♂ | 0.09 | 1.00 | 0.13 | <0.05 | 0.21 | <0.0001 | 0.21 | >0.01 | 0.44 | 1.00 | 0.40 | 1.00 |
| PWD ♀ vs. PWD ♂ | 0.29 | <0.0001 | 0.45 | <0.0001 | 0.37 | <0.0001 | 0.43 | >0.0001 | 0.46 | >0.01 | 0.49 | >0.001 |
| B6 ♀ vs. PWD ♀ | 0.21 | <0.0001 | 0.15 | <0.001 | 0.19 | <0.01 | 0.10 | 1.00 | 0.37 | 0.53 | 0.28 | 1.00 |
| B6 ♀ vs. 129 ♀ | 0.14 | <0.001 | 0.07 | 1.00 | 0.17 | <0.05 | 0.11 | 1.00 | 0.39 | 1.00 | 0.30 | 1.00 |
| B6 ♀ vs. CAST ♀ | 0.09 | 0.66 | 0.06 | 1.00 | 0.17 | 0.10 | 0.07 | 1.00 | 0.09 | 1.00 | 0.26 | 1.00 |
| B6 ♀ vs. DBA ♀ | 0.17 | <0.001 | 0.14 | <0.01 | 0.12 | 1.00 | 0.17 | 0.22 | 0.42 | >0.05 | 0.44 | >0.05 |
| PWD ♀ vs. 129 ♀ | 0.13 | 0.08 | 0.19 | <0.0001 | 0.12 | 1.00 | 0.17 | 0.20 | 0.16 | 1.00 | 0.15 | 1.00 |
| PWD ♀ vs. CAST ♀ | 0.22 | <0.0001 | 0.12 | 0.13 | 0.29 | <0.0001 | 0.13 | 1.00 | 0.33 | 1.00 | 0.20 | 1.00 |
| PWD ♀ vs. DBA ♀ | 0.15 | <0.05 | 0.13 | 0.20 | 0.27 | <0.001 | 0.23 | >0.05 | 0.17 | 1.00 | 0.19 | 1.00 |
| 129 ♀ vs. CAST ♀ | 0.19 | <0.0001 | 0.12 | 0.10 | 0.30 | <0.0001 | 0.13 | 1.00 | 0.33 | 1.00 | 0.20 | 1.00 |
| 129 ♀ vs. DBA ♀ | 0.13 | 0.11 | 0.17 | <0.01 | 0.25 | <0.0001 | 0.13 | 1.00 | 0.24 | 1.00 | 0.23 | 1.00 |
| CAST ♀ vs. DBA ♀ | 0.16 | <0.01 | 0.12 | 0.35 | 0.09 | 1.00 | 0.12 | 1.00 | 0.37 | 0.18 | 0.27 | 1.00 |
| DBA ♂ vs. 129 ♂ | 0.13 | 0.93 | 0.13 | 1.00 | 0.21 | 0.79 | 0.22 | 0.34 | 0.72 | 0.85 | 0.61 | 1.00 |
| DBA ♂ vs. PWD ♂ | 0.14 | 0.30 | 0.13 | 0.58000 | 0.11 | 1.00 | 0.08 | 1.00 | 0.76 | 0.11 | 0.72 | 0.23 |
| DBA ♂ vs. B6 ♂ | 0.14 | 0.25 | 0.14 | 0.30000 | 0.06 | 1.00 | 0.07 | 1.00 | 0.53 | 1.00 | 0.57 | 1.00 |
| DBA ♂ vs. CAST ♂ | 0.17 | 0.67 | 0.10 | 1.00 | 0.19 | 1.00 | 0.13 | 1.00 | 0.40 | 1.00 | 0.40 | 1.00 |
| 129 ♂ vs. PWD ♂ | 0.16 | <0.0001 | 0.16 | <0.0001 | 0.25 | <0.0001 | 0.18 | >0.01 | 0.19 | 1.00 | 0.20 | 1.00 |
| 129 ♂ vs. B6 ♂ | 0.16 | <0.0001 | 0.17 | <0.0001 | 0.20 | <0.0001 | 0.19 | >0.01 | 0.19 | 1.00 | 0.16 | 1.00 |
| 129 ♂ vs. CAST ♂ | 0.19 | <0.01 | 0.15 | 0.29 | 0.07 | 1.00 | 0.16 | 1.00 | 0.49 | 1.00 | 0.43 | 1.00 |
| PWD ♂ vs. B6 ♂ | 0.04 | 1.00 | 0.03 | 1.00 | 0.10 | 1.00 | 0.06 | 1.00 | 0.24 | 1.00 | 0.20 | 1.00 |
| PWD ♂ vs. CAST ♂ | 0.08 | 1.00 | 0.06 | 1.00 | 0.23 | <0.0001 | 0.12 | 1.00 | 0.54 | 1.00 | 0.51 | 1.00 |
| B6 ♂ vs. CAST ♂ | 0.06 | 1.00 | 0.07 | 1.00 | 0.19 | 0.23 | 0.11 | 1.00 | 0.47 | 1.00 | 0.37 | 1.00 |
| DBA ♀ vs. CAST ♂ | 0.24 | <0.001 | 0.31 | <0.0001 | 0.20 | 0.66 | 0.28 | >0.01 | 0.33 | 1.00 | 0.37 | 1.00 |
| DBA ♀ vs. B6 ♂ | 0.23 | <0.0001 | 0.36 | <0.0001 | 0.17 | 0.24 | 0.31 | >0.0001 | 0.42 | 0.17 | 0.50 | >0.01 |
| DBA ♀ vs. 129 ♂ | 0.20 | <0.0001 | 0.25 | <0.0001 | 0.17 | 0.49 | 0.23 | >0.01 | 0.49 | 0.15 | 0.52 | 0.06 |
| DBA ♀ vs. PWD ♂ | 0.23 | <0.0001 | 0.35 | <0.0001 | 0.19 | <0.05 | 0.27 | >0.0001 | 0.52 | >0.0001 | 0.65 | >0.0001 |
| CAST ♀ vs. DBA ♂ | 0.08 | 1.00 | 0.26 | <0.0001 | 0.15 | 1.00 | 0.39 | >0.0001 | 0.61 | 1.00 | 0.43 | 1.00 |
| CAST ♀ vs. B6 ♂ | 0.09 | 1.00 | 0.36 | <0.0001 | 0.13 | 1.00 | 0.38 | >0.0001 | 0.11 | 1.00 | 0.37 | 0.38 |
| CAST ♀ vs. 129 ♂ | 0.15 | <0.001 | 0.22 | <0.0001 | 0.22 | <0.01 | 0.27 | >0.0001 | 0.16 | 1.00 | 0.43 | 0.42 |
| CAST ♀ vs. PWD ♂ | 0.09 | 0.36 | 0.34 | <0.0001 | 0.10 | 1.00 | 0.34 | >0.0001 | 0.19 | 1.00 | 0.46 | >0.0001 |
| B6 ♀ vs. DBA ♂ | 0.07 | 1.00 | 0.27 | <0.0001 | 0.19 | 1.00 | 0.41 | >0.0001 | 0.67 | 0.74 | 0.46 | 1.00 |
| B6 ♀ vs. CAST ♂ | 0.12 | 1.00 | 0.30 | <0.0001 | 0.13 | 1.00 | 0.36 | >0.0001 | 0.40 | 1.00 | 0.38 | 1.00 |
| B6 ♀ vs. 129 ♂ | 0.09 | 0.16 | 0.18 | <0.0001 | 0.08 | 1.00 | 0.28 | >0.0001 | 0.13 | 1.00 | 0.38 | 1.00 |
| B6 ♀ vs. PWD ♂ | 0.08 | <0.05 | 0.32 | <0.0001 | 0.21 | <0.0001 | 0.37 | >0.0001 | 0.15 | 1.00 | 0.29 | 0.24 |
| 129 ♀ vs. DBA ♂ | 0.16 | 0.18 | 0.24 | <0.0001 | 0.33 | <0.0001 | 0.37 | >0.0001 | 0.36 | 1.00 | 0.44 | 1.00 |
| 129 ♀ vs. CAST ♂ | 0.23 | <0.001 | 0.27 | <0.0001 | 0.23 | <0.05 | 0.31 | >0.0001 | 0.29 | 1.00 | 0.29 | 1.00 |
| 129 ♀ vs. B6 ♂ | 0.22 | <0.0001 | 0.29 | <0.0001 | 0.35 | <0.0001 | 0.34 | >0.0001 | 0.30 | 1.00 | 0.38 | 1.00 |
| 129 ♀ vs. PWD ♂ | 0.22 | <0.0001 | 0.28 | <0.0001 | 0.36 | <0.0001 | 0.33 | >0.0001 | 0.47 | 0.19 | 0.49 | 0.10 |
| PWD ♀ vs. DBA ♂ | 0.19 | <0.05 | 0.35 | <0.0001 | 0.33 | <0.0001 | 0.49 | >0.0001 | 0.38 | 1.00 | 0.46 | 1.00 |
| PWD ♀ vs. CAST ♂ | 0.34 | <0.0001 | 0.42 | <0.0001 | 0.30 | <0.0001 | 0.43 | >0.0001 | 0.36 | 1.00 | 0.33 | 1.00 |
| PWD ♀ vs. B6 ♂ | 0.29 | <0.0001 | 0.46 | <0.0001 | 0.34 | <0.0001 | 0.47 | >0.0001 | 0.32 | 1.00 | 0.36 | 1.00 |
| PWD ♀ vs. 129 ♂ | 0.18 | <0.0001 | 0.31 | <0.0001 | 0.24 | <0.0001 | 0.36 | >0.0001 | 0.42 | 1.00 | 0.44 | 0.85 |
