## Supplementary material for "Chromosome length is not the sole determinant of sexually dimorphic crossover rates during mammalian meiosis: Insights from genetically diverse mouse strains": Table S4

| | MLH1 foci per SC | | Median distance from centromere to first MLH1 focus | | | | | | | | | |
|---|---|---|---|---|---|---|---|---|---|---|---|---|
| | | | DBA/2J ♂ | DBA/2J ♀ | CAST/EiJ ♂ | CAST/EiJ ♀ | C57Bl/6J ♂ | C57Bl/6J ♀ | 129S1/SvImJ ♂ | 129S1/SvImJ ♀ | PWD/PhJ ♂ | PWD/PhJ ♀ |
| All SCs | 1 | Absolute distance (µm) | 4.84 | 4.64 | 4.83 | 4.67 | 5.06 | 4.70 | 4.73 | 4.20 | 4.83 | 4.70 |
| | | Normalized distance (%SC length) | 69.49 | 53.63 | 69.29 | 54.13 | 70.22 | 53.77 | 66.22 | 56.48 | 74.06 | 56.97 |
| | >1 | Absolute distance (µm) | 2.10 | 2.53 | 2.53 | 2.60 | 2.24 | 2.72 | 2.46 | 2.36 | 2.09 | 2.97 |
| | | Normalized distance (%SC length) | 23.32 | 23.70 | 25.33 | 23.80 | 24.14 | 25.32 | 26.33 | 24.74 | 21.60 | 29.43 |

| | SCs with 1 MLH1 focus | | | | SCs with >1 MLH1 focus | | | |
|---|---|---|---|---|---|---|---|---|
| | Absolute Distance (µm) | | Normalized Distance (%SC Length) | | Absolute Distance (µm) | | Normalized Distance (%SC Length) | |
| Comaprison | KS Stat | Adj. P value | KS Stat | Adj.P value | KS Stat | Adj. P value | KS Stat | Adj.P value |
| DBA ♀ vs. DBA ♂ | 0.05 | 1.00 | 0.21 | <0.0001 | 0.20 | <0.05 | 0.17 | 0.13 |
| CAST ♀ vs. CAST ♂ | 0.08 | 1.00 | 0.26 | <0.0001 | 0.11 | 1.00 | 0.09 | 1.00 |
| B6 ♀ vs. B6 ♂ | 0.07 | 0.05 | 0.27 | <0.0001 | 0.18 | <0.0001 | 0.09 | 0.11 |
| 129 ♀ vs. 129 ♂ | 0.13 | <0.0001 | 0.17 | <0.0001 | 0.04 | 1.00 | 0.09 | 1.00 |
| PWD ♀ vs. PWD ♂ | 0.06 | 1.00 | 0.27 | <0.0001 | 0.29 | <0.0001 | 0.26 | <0.0001 |
| B6 ♀ vs. PWD ♀ | 0.04 | 1.00 | 0.06 | 0.90 | 0.09 | 0.65 | 0.15 | <0.001 |
| B6 ♀ vs. 129 ♀ | 0.10 | <0.001 | 0.06 | 1.00 | 0.16 | <0.0001 | 0.07 | 1.00 |
| B6 ♀ vs. CAST ♀ | 0.03 | 1.00 | 0.03 | 1.00 | 0.06 | 1.00 | 0.08 | 1.00 |
| B6 ♀ vs. DBA ♀ | 0.05 | 1.00 | 0.07 | 0.47 | 0.12 | 0.10 | 0.13 | <0.05 |
| PWD ♀ vs. 129 ♀ | 0.10 | <0.01 | 0.04 | 1.00 | 0.20 | <0.0001 | 0.19 | <0.0001 |
| PWD ♀ vs. CAST ♀ | 0.06 | 1.00 | 0.06 | 1.00 | 0.13 | <0.05 | 0.20 | <0.0001 |
| PWD ♀ vs. DBA ♀ | 0.08 | 0.11 | 0.09 | >0.01 | 0.16 | <0.01 | 0.22 | <0.0001 |
| 129 ♀ vs. CAST ♀ | 0.12 | <0.01 | 0.06 | 1.00 | 0.13 | 0.08 | 0.06 | 1.00 |
| 129 ♀ vs. DBA ♀ | 0.11 | <0.0001 | 0.06 | 1.00 | 0.10 | 1.00 | 0.09 | 1.00 |
| CAST ♀ vs. DBA ♀ | 0.05 | 1.00 | 0.06 | 1.00 | 0.07 | 1.00 | 0.12 | 0.39 |
| DBA ♂ vs. 129 ♂ | 0.09 | <0.01 | 0.08 | <0.05 | 0.14 | 0.51 | 0.14 | 0.46 |
| DBA ♂ vs. PWD ♂ | 0.06 | 1.00 | 0.08 | <0.0001 | 0.06 | 1.00 | 0.10 | 1.00 |
| DBA ♂ vs. B6 ♂ | 0.05 | 0.45 | 0.08 | <0.0001 | 0.07 | 1.00 | 0.07 | 1.00 |
| DBA ♂ vs. CAST ♂ | 0.09 | <0.05 | 0.07 | 1.00 | 0.18 | 0.53 | 0.13 | 1.00 |
| 129 ♂ vs. PWD ♂ | 0.05 | 1.00 | 0.15 | <0.0001 | 0.14 | <0.0001 | 0.15 | <0.0001 |
| 129 ♂ vs. B6 ♂ | 0.09 | <0.001 | 0.11 | <0.0001 | 0.08 | 0.77 | 0.09 | 0.22 |
| 129 ♂ vs. CAST ♂ | 0.04 | 1.00 | 0.10 | <0.01 | 0.05 | 1.00 | 0.05 | 1.00 |
| PWD ♂ vs. B6 ♂ | 0.06 | 0.12 | 0.06 | 0.16 | 0.08 | 0.06 | 0.09 | 0.01 |
| PWD ♂ vs. CAST ♂ | 0.04 | 1.00 | 0.07 | 0.39 | 0.16 | <0.05 | 0.15 | 0.15 |
| B6 ♂ vs. CAST ♂ | 0.06 | 1.00 | 0.05 | 1.00 | 0.11 | 1.00 | 0.10 | 1.00 |
| DBA ♀ vs. CAST ♂ | 0.11 | <0.001 | 0.27 | <0.0001 | 0.11 | 1.00 | 0.14 | 1.00 |
| DBA ♀ vs. B6 ♂ | 0.09 | <0.0001 | 0.27 | <0.0001 | 0.13 | <0.05 | 0.14 | >0.05 |
| DBA ♀ vs. 129 ♂ | 0.10 | <0.01 | 0.21 | <0.0001 | 0.10 | 1.00 | 0.14 | >0.05 |
| DBA ♀ vs. PWD ♂ | 0.08 | <0.05 | 0.33 | <0.0001 | 0.15 | <0.001 | 0.13 | >0.01 |
| CAST ♀ vs. DBA ♂ | 0.04 | 1.00 | 0.22 | <0.0001 | 0.22 | <0.0001 | 0.09 | 1.00 |
| CAST ♀ vs. B6 ♂ | 0.08 | 0.74 | 0.23 | <0.0001 | 0.16 | <0.0001 | 0.05 | 1.00 |
| CAST ♀ vs. 129 ♂ | 0.10 | 0.26 | 0.20 | <0.0001 | 0.12 | <0.05 | 0.10 | 0.48 |
| CAST ♀ vs. PWD ♂ | 0.06 | 1.00 | 0.27 | <0.0001 | 0.18 | <0.0001 | 0.08 | 1.00 |
| B6 ♀ vs. DBA ♂ | 0.12 | <0.0001 | 0.26 | <0.0001 | 0.24 | <0.0001 | 0.14 | 0.41 |
| B6 ♀ vs. CAST ♂ | 0.14 | <0.0001 | 0.25 | <0.0001 | 0.12 | 1.00 | 0.05 | 1.00 |
| B6 ♀ vs. 129 ♂ | 0.16 | <0.0001 | 0.22 | <0.0001 | 0.14 | <0.0001 | 0.04 | 1.00 |
| B6 ♀ vs. PWD ♂ | 0.14 | <0.0001 | 0.32 | <0.0001 | 0.21 | <0.0001 | 0.14 | >0.0001 |
| 129 ♀ vs. DBA ♂ | 0.03 | 1.00 | 0.21 | <0.0001 | 0.12 | 1.00 | 0.13 | 1.00 |
| 129 ♀ vs. CAST ♂ | 0.07 | 0.74 | 0.22 | <0.0001 | 0.07 | 1.00 | 0.07 | 1.00 |
| 129 ♀ vs. B6 ♂ | 0.08 | 0.07 | 0.22 | <0.0001 | 0.06 | 1.00 | 0.08 | 1.00 |
| 129 ♀ vs. PWD ♂ | 0.05 | 1.00 | 0.28 | <0.0001 | 0.11 | <0.05 | 0.12 | >0.01 |
| PWD ♀ vs. DBA ♂ | 0.05 | 1.00 | 0.21 | <0.0001 | 0.31 | <0.0001 | 0.23 | >0.001 |
| PWD ♀ vs. CAST ♂ | 0.07 | 1.00 | 0.21 | <0.0001 | 0.18 | 0.08 | 0.15 | 0.49 |
| PWD ♀ vs. B6 ♂ | 0.09 | <0.01 | 0.22 | <0.0001 | 0.24 | <0.0001 | 0.19 | >0.0001 |
| PWD ♀ vs. 129 ♂ | 0.07 | 0.49 | 0.16 | <0.0001 | 0.19 | <0.0001 | 0.12 | >0.05 |
