## Supplementary material for "Chromosome length is not the sole determinant of sexually dimorphic crossover rates during mammalian meiosis: Insights from genetically diverse mouse strains": Table S5

| **Antibody** | **Dilution Used** | **Vendor** | **Catalog Number** |
|---|---|---|---|
| Mouse anti-MLH1 mAb | 1:100 | BD Biosciences | 550838 |
| Guinea pig anti-MLH3 pAb | 1:500 | Custom made with Thermo-Fisher | Horan et al., 2024 |
| Rabbit anti-RAD51 pAb | 1:500 | Millipore | PC130 |
| Rabbit anti-MSH4 pAb | 1:100 | ABclonal | A8556 |
| Rabbit anti-CCNB1IP1 | 1:100 | ABclonal | A16693 |
| Mouse anti-SYCP3 mAb | 1:1,000 | Abcam | ab97672 |
| Rabbit anti-SYCP3 pAb | 1:10,000 | Custome made | Kolas et al., 2005 |
| Human anti-Centromere Protein pAb | 1:1,000 | Antibodies Inc. | 15-234 |
| Alexa Fluor® 488 AffiniPure® $F(ab')_2$ Fragment Goat Anti-Mouse IgG, Fcγ fragment specific | 1:1,000 | Jackson ImmunoResearch | 115-546-008 |
| Rhodamine Red™-X (RRX) AffiniPure® $F(ab')_2$ Fragment Goat Anti-Mouse IgG, Fcγ fragment specific | 1:1,000 | Jackson ImmunoResearch | 115-296-071 |
| Alexa Fluor® 488 AffiniPure® $F(ab')_2$ Fragment Goat Anti-Rabbit IgG, Fc fragment specific | 1:1,000 | Jackson ImmunoResearch | 111-546-046 |
| Rhodamine Red™-X (RRX) AffiniPure® $F(ab')_2$ Fragment Goat Anti-Rabbit IgG, Fc fragment specific | 1:1,000 | Jackson ImmunoResearch | 111-296-046 |
| Alexa Fluor® 647 AffiniPure® $F(ab')_2$ Fragment Goat Anti-Guinea Pig IgG (H+L) | 1:1,000 | Jackson ImmunoResearch | 106-606-003 |
| Alexa Fluor® 647 AffiniPure® $F(ab')_2$ Fragment Goat Anti-Human IgG, Fcγ fragment specific | 1:1,000 | Jackson ImmunoResearch | 109-606-170 |
